# MCB-613 exploits a collateral sensitivity in drug resistant *EGFR*-mutant non-small cell lung cancer through covalent inhibition of KEAP1

**DOI:** 10.1101/2023.01.17.524094

**Authors:** Christopher F. Bassil, Gray R. Anderson, Benjamin Mayro, Kayleigh N. Askin, Peter S. Winter, Samuel Gruber, Tierney M. Hall, Jacob P. Hoj, Christian Cerda-Smith, Haley M. Hutchinson, Shane T. Killarney, Katherine R. Singleton, Li Qin, Kévin Jubien-Girard, Cécile Favreau, Anthony R. Martin, Guillaume Robert, Rachid Benhida, Patrick Auberger, Ann Marie Pendergast, David M. Lonard, Alexandre Puissant, Kris C. Wood

## Abstract

Targeted therapies have revolutionized cancer chemotherapy. Unfortunately, most patients develop multifocal resistance to these drugs within a matter of months. Here, we used a high-throughput phenotypic small molecule screen to identify MCB-613 as a compound that selectively targets *EGFR*-mutant, EGFR inhibitor-resistant non-small cell lung cancer (NSCLC) cells harboring diverse resistance mechanisms. Subsequent proteomic and functional genomic screens involving MCB-613 identified its target in this context to be KEAP1, revealing that this gene is selectively essential in the setting of EGFR inhibitor resistance. In-depth molecular characterization demonstrated that (1) MCB-613 binds KEAP1 covalently; (2) a single molecule of MCB-613 is capable of bridging two KEAP1 monomers together; and, (3) this modification interferes with the degradation of canonical KEAP1 substrates such as NRF2. Surprisingly, NRF2 knockout sensitizes cells to MCB-613, suggesting that the drug functions through modulation of an alternative KEAP1 substrate. Together, these findings advance MCB-613 as a new tool for exploiting the selective essentiality of KEAP1 in drug-resistant, *EGFR*-mutant NSCLC cells.

## INTRODUCTION

Cancer is the second leading cause of death worldwide.^1^ Although the use of targeted therapies has resulted in encouraging clinical responses, these benefits are, unfortunately, often short-lived; in the case of non-small cell lung cancers (NSCLC) driven by activating mutations in the epidermal growth factor receptor (EGFR), for instance, most patients will develop resistance to first-line targeted therapy within 24 months.^2,3^ This problem is complicated by two interrelated considerations: (1) *EGFR*-mutant NSCLC cells can develop resistance to EGFR inhibitors through a wide variety of distinct mechanisms; and, (2) many, if not all, of these mechanisms can coevolve simultaneously in the same patient or tumor.^4^ This presents clinicians and researchers with a difficult quandary: on the one hand, treatments which use one drug to target an individual resistance mechanism are insufficient and unlikely to prove curative; on the other hand, treatments which use multiple drugs to target many different mechanisms at the same time are demanding, and are unlikely to be feasible. Thus, new approaches to the problem of “multifocal” drug resistance are needed.

One strategy is to identify, target, and exploit vulnerabilities which emerge as a consequence of drug resistance itself. These acquired, or “collateral,” sensitivities—which are distinct from chemical synthetic lethalities in that they persist even after removal of the original, selecting drug—have long been documented in the microbial literature.^5^ Recently, for instance, it was shown that clinical isolates taken from the lungs of cystic fibrosis patients suffering from chronic, drug-resistant *Pseudomonas aeruginosa* infections consistently harbored mutations in the *nfxB* gene.^6^ These mutations, though pathoadaptive in the setting of conventional anti-pseudomonal therapy with fluoroquinolones, nevertheless rendered these bacterial populations collaterally sensitive to the commonly used and readily available aminoglycoside antibiotic amikacin. In point of fact, drug resistance often promotes targetable collateral sensitivities to commonplace, existing drugs. In the case of the recently approved fluorocycline antibiotic eravacycline, for example, drug resistance in *Klebsiella pneumoniae* promotes sensitivity to commonly used antibiotics like aztreonam and ceftazidime.^7^

In recent years, this concept has increasingly been applied to the study of drug resistance in cancer as well. It has been demonstrated, for example, that the acquisition of BRAF inhibitor resistance in *BRAF*-mutant melanoma engenders an upregulation of reactive oxygen species (ROS) which in some cells promotes a stable and clinically relevant collateral sensitivity to histone deacetylase (HDAC) inhibitors.^8^ Separately, previous work from our group has shown that diverse pathways of drug resistance in this setting also converge upon the activation of the c-MYC transcription factor; this, in turn, gives rise to shared, actionable collateral sensitivities to drugs which target MYC synthetic lethal partners such as certain tyrosine kinase families and metabolic pathways.^9^ Since at least some of these targetable collateral sensitivities to cancer drug resistance are both stable and predictable, moreover, it is conceivable that future first-line therapies could be designed specifically to guide tumor evolution toward targetable, collaterally sensitive states. Indeed, previous work from our group has shown that BET bromodomain inhibitors can be used to guide acute myeloid leukemia (AML) cells into an “evolutionary trap” which collaterally sensitizes them to existing, clinically approved BCL2 inhibitors.^10^ Finally, these effects are legion: with examples cited in a growing number of cases, it appears increasingly likely that numerous such collateral sensitivities may in fact be hidden throughout the landscape of drug-resistant cancers.^11–15^

Here, we report on the discovery of a collateral sensitivity shared across multiple, distinct mechanisms of EGFR inhibitor resistance in *EGFR*-mutant non-small cell lung cancer (NSCLC). Specifically, we show that the acquisition of EGFR inhibitor resistance by diverse mechanisms sensitizes *EGFR*-mutant NSCLC cells to a small molecule known as MCB-613. This effect relies upon the covalent destabilization of Kelch-like ECH associated protein 1 (KEAP1). More precisely, this electrophilic small molecule uses two distinct Michael acceptor sites to tether monomers of KEAP1 together. The perturbation of KEAP1 by MCB-613 leads to the dissociation of KEAP1 from its canonical repression substrate, nuclear factor erythroid 2-related factor 2 (NRF2), resulting in the stabilization of NRF2 protein and the downstream activation of its transcriptional program. These effects, however, are dispensable for the selective cell death mechanism of MCB-613, raising the intriguing possibility that it is the accumulation of an alternative—and perhaps heretofore undescribed—substrate of KEAP1 that is responsible for this shared collateral sensitivity.

## RESULTS

### A pharmacologic screen to identify shared collateral sensitivities in *EGFR*-mutant, EGFR inhibitor-resistant NSCLC

To identify targetable collateral sensitivities shared across distinct mechanisms of EGFR inhibitor resistance, we compiled a panel of four well-characterized *EGFR*-mutant non-small cell lung cancer cell lines with varying degrees of resistance to EGFR inhibition: PC9, PFR3, GR4, and WZR12. The “parental” PC9 cell line is drug-naïve and retains exquisite sensitivity to EGFR inhibition. The previously characterized PFR3, GR4, and WZR12 cell lines—which are all derived from PC9s and driven by diverse, clinically relevant resistance mechanisms including IGF1R activation, *EGFR* T790M secondary site mutation, and downstream *MAPK1* amplification, respectively—display increasing degrees of resistance to the first-generation EGFR inhibitor gefinitib (Fig. 1a).^16–18^ We screened all four of these cell lines against a library of 2,100 small molecules in a three-day viability assay at two separate drug doses (Fig. 1b). As expected, we detected numerous collateral resistances and sensitivities to EGFR inhibitor resistance (Fig. 1c). Predictably, receptor tyrosine kinase inhibitors, including numerous other EGFR inhibitors (gefitinib, erlotinib, and afatinib, among others), scored as shared collateral resistances, providing an important internal positive control. Of the collateral sensitivities, however, one in particular stood out across all three drug-resistant derivatives: a relatively understudied molecule called MCB-613 (Fig. 1d). A follow-up, three-day, eight-point GI_50_ assay confirmed that the three drug-resistant derivatives were collaterally sensitive to MCB-613, and further revealed that there was a strong, direct, stepwise correlation between the degrees of resistance to EGFR inhibition on the one hand and sensitivity to MCB-613 on the other (R^2^ = 0.9962) (Fig. 1e-f).

**Fig. 1.**
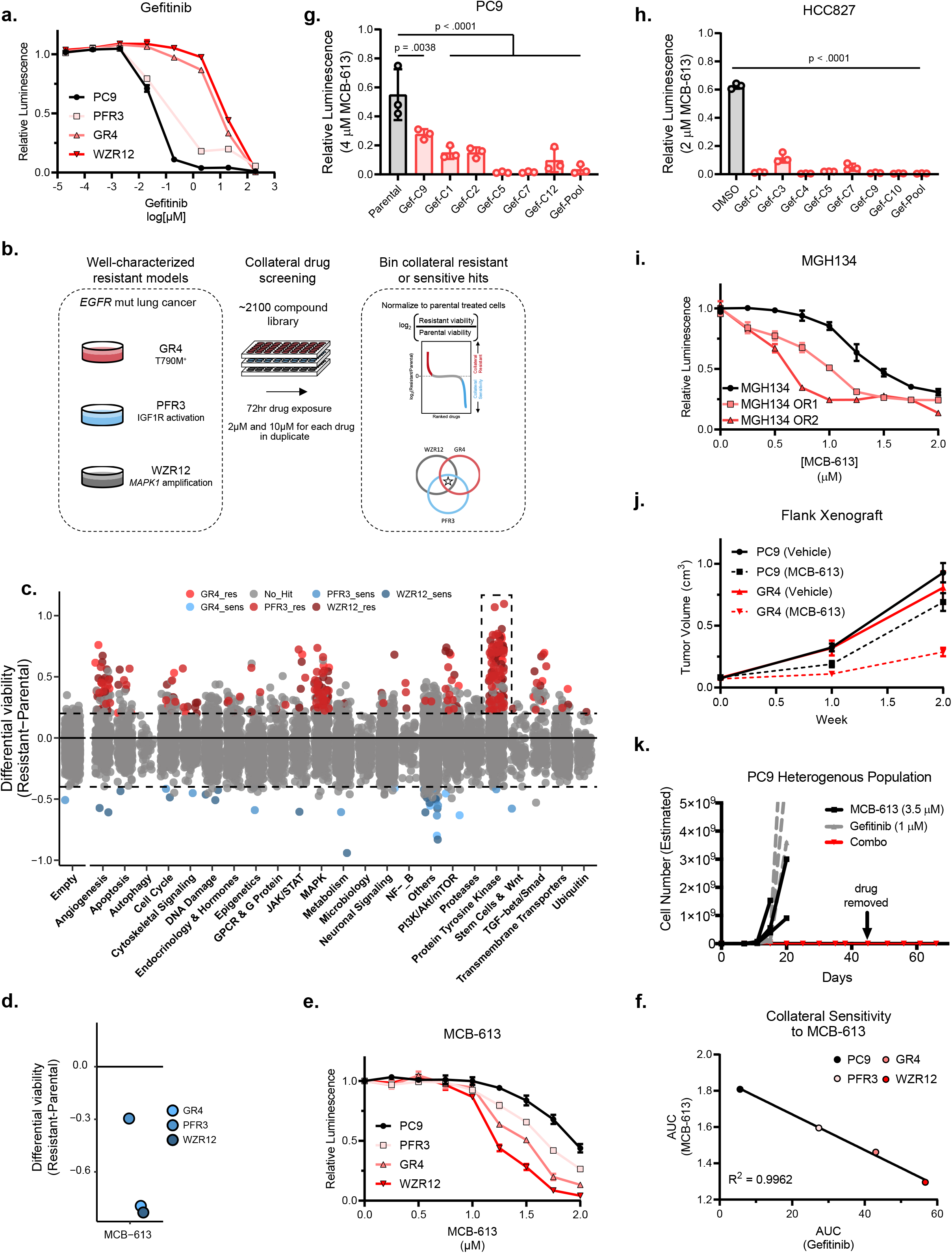
Drug resistance promotes collateral sensitivity to MCB-613 in *EGFR-mutant* NSCLC. a. Relative cell viability of parental (PC9) and diverse drug-resistant (PFR3, GR4, WZR12) *EGFR-mutant* NSCLC lines following 72-hour incubation with gefitinib across an 8-point serial drug dilution series. Data are mean ± SEM for n = 3 biologically independent experiments. b. High-throughput pharmacologic screening strategy to identify collateral effects across a panel of parental (not shown) and drug-resistant cells harboring diverse mechanisms of resistance. c. Manhattan plot depicting results of chemical screen. Each point represents the average of log2(resistant/parental) from two biologically independent experiments. Collateral resistances and sensitivities are shown in red and blue, respectively. Gray indicates all other conditions (no collateral effect, empty control wells, vehicle (DMSO) control wells). Dotted box indicates widespread resistance to other receptor tyrosine kinase inhibitors as a positive control. d. Magnification of the Manhattan plot depicted in (c) highlighting MCB-613. e. Relative cell viability of parental (PC9) and diverse drug-resistant (PFR3, GR4, WZR12) cells following 72-hour incubation with MCB-613 across an 8-point linear drug dilution series. Data are mean ± SEM for n = 3 biologically independent experiments. f. Scatterplot depicting correlation between area-under-the-curve (AUC) calculations for GI_50_ dose-response curves for gefitinib (x-axis) depicted in (a) and MCB-613 (y-axis) depicted in (e). g. Histogram depicting cell viability of parental PC9 cells, a gefitinib-resistant PC9 pooled population, and six gefitinib-resistant PC9 clonal populations after 72 hours of exposure to a 4 μM dose of MCB-613. Data are mean ± SEM for n = 3 biologically independent experiments. h. Histogram depicting cell viability of parental HCC827 cells, the previously described gefitinib-resistant HCC827 GR6 clone, and seven freshly derived gefitinib-resistant HCC827 clonal populations after 72 hours of exposure to a 2 μM dose of MCB-613. Data are mean ± SEM of n = 3 biologically independent experiments. i. Relative cell viability of matched parental (MGH134) and osimertinib-resistant (MGH134 OR1 and OR2) patient-derived cell lines following 72-hour incubation with MCB-613 across an 8-point linear drug dilution series. Data are mean ± SEM for n = 3 biologically independent experiments. j. Tumor growth curves from 5-6 week-old female SCID mice xenotransplanted with parental PC9 or drug-resistant GR4 cells and treated with vehicle (saline) or MCB-613 (20 mg/kg) three times weekly via interaperitoneal injection. Data are mean ± SEM for n = 10 biologically independent experiments. k. Line plot depicting estimated cumulative cell growth over time for a heterogeneous population of drug-naïve and drug-resistant PC9 cells exposed to either gefitinib, MCB-613, or a combination of the two. Data are mean ± SEM for n = 3 biologically independent experiments.

We next sought to assess the breadth of this shared collateral sensitivity to EGFR inhibitor resistance. First, we gathered together two drug-naïve, *EGFR*-mutant NSCLC cell lines—PC9 and HCC827—which are known to develop EGFR inhibitor resistance through diverse mechanisms.^19–22^ We used increasing doses of gefitinib to cultivate EGFR inhibitor resistance in these cell lines before selecting individual clones and querying them for sensitivity to MCB-613 (Supplementary Fig. 1a-b). In all clones tested, resistance to EGFR inhibition was associated with collateral sensitivity to MCB-613 (Fig. 1g-h). Finally, to assess whether this finding was limited to gefitinib or extended to other clinically relevant EGFR inhibitors, we evolved MGH134, a previously described, patient-derived cell line model of *EGFR*-mutant NSCLC, resistant to the first-line, third-generation small molecule inhibitor osimertinib; in accordance with the above findings, resistance to this standard-of-care targeted therapy also conferred increased sensitivity to MCB-613 (Fig. 1i, Supplementary Fig. 1c).^23^

Lastly, to evaluate whether the broad, shared collateral sensitivity of *EGFR*-mutant, EGFR inhibitor-resistant NSCLC cells to MCB-613 represents a viable preclinical strategy for overcoming multifocal drug resistance, we used a flank xenograft model to implant either drug-naïve PC9 or drug-resistant GR4 cells into SCID mice and treated them with MCB-613. As expected, the drug-resistant GR4 derivatives were selectively suppressed by MCB-613 (Fig. 1j). Given the strong correlation between resistance to EGFR inhibition and sensitivity to MCB-613 observed above, we wondered whether the combination of these two therapeutic approaches in a single *EGFR*-mutant NSCLC population would suffice to successfully suppress all cellular outgrowth. To test this hypothesis, we mixed drug-naïve and drug-resistant PC9 cells together and then exposed this heterogeneous population to either gefitinib alone, MCB-613 alone, or a combination of the two. Although neither molecule on its own suppressed outgrowth, the combination of the two was enough to completely eradicate the population (Fig. 1k). When considered together, these findings establish the widespread therapeutic relevance of MCB-613 as a *bona fide* collateral sensitivity shared across distinct mechanisms of resistance to EGFR inhibition.

### A click-chemistry approach to mapping the interactome of MCB-613

Phenotypic drug discovery screens such as the one described above are useful for finding biologically active compounds; they also, however, frequently identify lesser studied, promiscuous molecules that lack specific, well-characterized targets.^24^ As a sparsely studied molecule itself, MCB-613 (4-ethyl-2,6-bis-pyridin-3-ylmethylene-cyclohexanone) presented us with several challenges to follow-up target identification and validation. First, MCB-613 has previously been shown to influence numerous, diverse cellular pathways; as such, the molecular target responsible for the collateral sensitivity observed above was not immediately apparent to us.^25–27^ Furthermore, the highly reactive structure of the molecule—which bears a central carbonyl flanked by two electrophilic, α, β-unsaturated bonds—provides ample opportunity for promiscuous interaction with a wide range of nucleophilic protein targets, possibly through a double Michael addition (Fig. 2a-b).^28^ Thus, a comprehensive strategy for mapping the protein interactome of MCB-613 was needed.

**Fig. 2.**
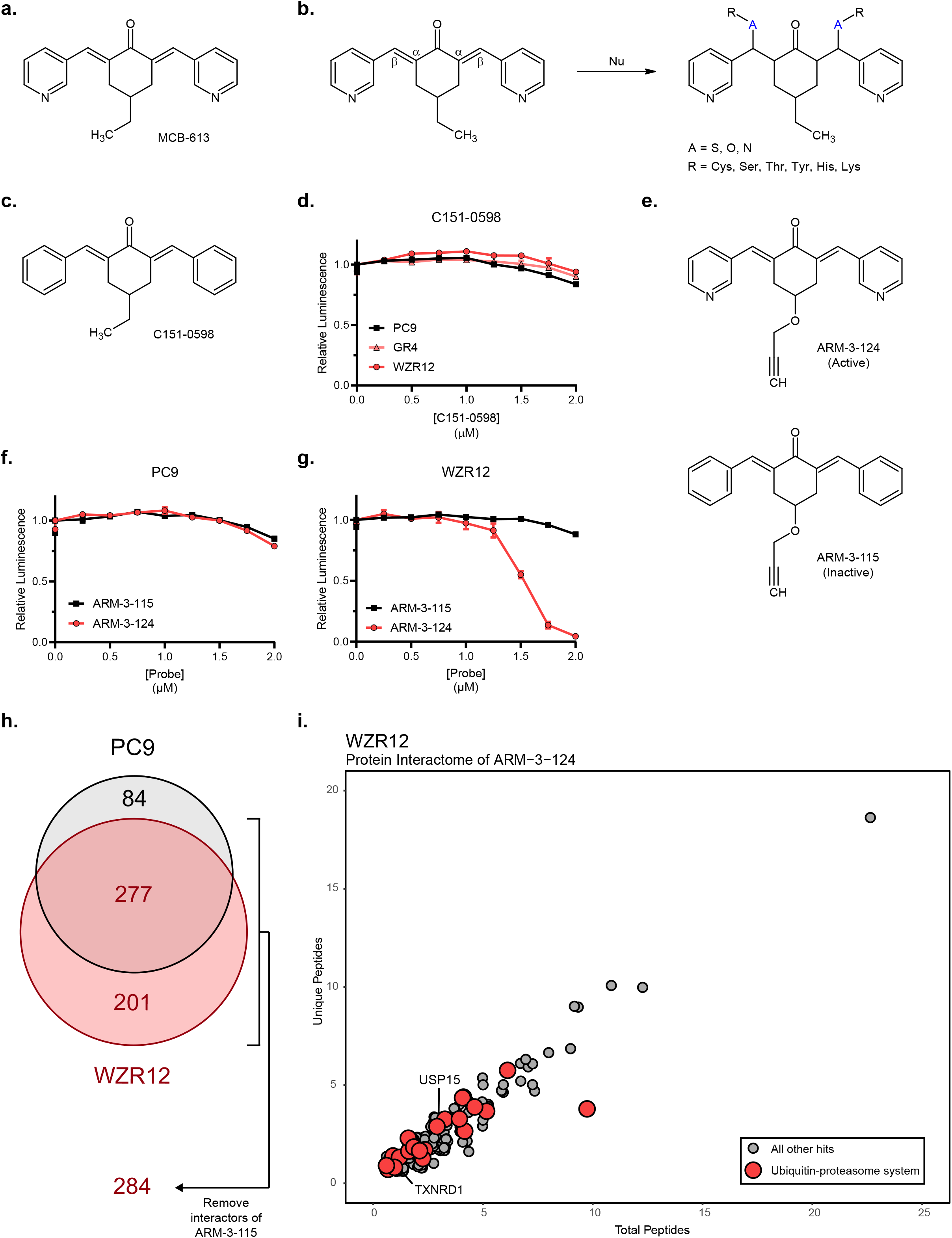
Structure of MCB-613 and clickable derivatives for mapping the protein interactome. a. Chemical structure of MCB-613. b. Diagram showing proposed double Michael addition between MCB-613 and nucleophilic amino acids. c. Chemical structure of C151-0598. d. Relative cell viability of parental (PC9) and diverse drug-resistant (GR4, WZR12) cells following 72-hour incubation with C151-0598 across an 8-point linear drug dilution series. Data are mean ± SEM for n = 3 biologically independent experiments. e. Chemical structures of the active ARM-3-124 (top) and inactive ARM-3-115 (bottom) clickable derivatives. f. Relative cell viability of parental PC9 cells following 72-hour incubation with the active ARM-3-124 (red) and inactive ARM-3-115 (black) clickable derivatives across an 8-point linear drug dilution series. Data are mean ± SEM for n = 3 biologically independent experiments. g. Relative cell viability of drug-resistant WZR12 cells following 72-hour incubation with the active ARM-3-124 (red) and inactive ARM-3-115 (black) clickable derivatives across an 8-point linear drug dilution series. Data are mean ± SEM for n = 3 biologically independent experiments. h. Venn-diagram illustrating results of click study. 84 proteins identified by ARM-3124 in the parental PC9 cells alone are depicted in black. 201 proteins identified by ARM-3-124 in the drug-resistant WZR12 cells alone are depicted in red. 277 proteins identified by ARM-3-124 in both parental and drug-resistant cells are depicted by the overlap. Removal from the 478 hits in the latter two groups of proteins which also bound the inactive ARM-3-115 yields a final list of 284. i. Scatterplot depicting protein interactors of ARM-3-124 in drug-resistant WZR12 cells. Large red circles indicate hits related to the ubiquitin-proteasome system. Labeled points represent known interactors of MCB-613.

Using C151-0598, a molecule which bears close structural similarity to MCB-613 but shows severely attenuated activity, we were able to design and implement a “click chemistry,” azide-alkyne cycloaddition (CuAAC) based approach to differential affinity purification and target identification (Fig. 2c-d). First, we generated “clickable” derivatives of MCB-613 and C151-0598 by replacing the ethylene units at the 4-positions of their central cyclohexanone moieties with propargyl ethers (Fig. 2e, Supplementary Methods). We called these active and inactive clickable derivatives “ARM-3-124” and “ARM-3-115,” respectively. After validating that the derivatives phenocopied the effects of their parent compounds in drug-naïve and drug-resistant *EGFR*-mutant NSCLC cells, we deployed them for target identification (Fig. 2f-g). Specifically, we exposed parental PC9 cells and their drug-resistant WZR12 derivatives to a dose which only showed efficacy in the ARM-3-124/WZR12 combination (2 μM) for one hour before collection, cell lysis, and copper-catalyzed azide-alkyne cycloaddition. These samples were then analyzed by mass spectrometry to identify bound peptides.

Across all four conditions, this approach identified >1,000 unique peptides corresponding to 610 protein interactors. After removing proteins which scored only in the inactive ARM-3-115 negative control conditions, this list was reduced to 560 proteins which scored with ARM-3-124 (regardless of ARM-3-115); after further removing proteins which scored only in the parental PC9 cells (reasoning that these would be less likely to cause selective cell death in the drug-resistant derivatives) the list was further refined to a total of 478 putative interactors. Although we carried all 478 of these interactors forward, we also paid special attention to the 284 proteins which scored as interactors of the active ARM-3-124 derivative but not the inactive ARM-3-115.

Further analysis yielded several interesting insights. First, the interactomes of the active derivative, ARM-3-124, demonstrated substantial overlap between the drug-naïve cells and drug-resistant derivatives, with a total of 277 hits (58% of total) shared between the two conditions (Fig. 2h). Second, a substantial number of proteins related to the ubiquitin-proteasome system (UPS) were also identified, consistent with recent reports that α, β-unsaturated compounds such as MCB-613 tend to show a predilection for such substrates. (This was consistent with the presence at low levels of thioredoxin reductase 1 (TXNRD1), another known substrate of the chalcone family of molecules to which MCB-613 belongs, as well as UPS15, a previously described substrate of MCB-613 in particular (Fig. 2i).)^26,27^ Together, these findings define a network of proteins which may be enriched for the functional target(s) of MCB-613 in EGFR inhibitor-resistant NSCLC cells.

### A CRISPR/Cas9 loss-of-function screen nominates KEAP1 as the molecular target of MCB-613

Having nominated a long list of potential targets, we next sought to rapidly and quantitatively evaluate the potential functional contributions of each of these 478 protein interactors to the collateral sensitivity of EGFR inhibitor-resistant NSCLC cells to MCB-613. Thus, we designed a targeted, loss-of-function CRISPR/Cas9 single-guide RNA (sgRNA) library containing a total of 3,344 sgRNAs targeting the genes encoding for each of the 478 interactors identified above. (Also included were 408 sgRNAs targeting 51 positive control genes encoding for ribosomal proteins, as well as 100 “safe harbor” guides, which target the human *AAVS1* locus and reliably produce no discernible phenotype.)^29^ We then transduced the parental PC9 and drug-resistant GR4 and WZR12 cell lines at a low multiplicity of infection (MOI) and cultured them under steady-state conditions for approximately 14 population doublings before de-convoluting the initial and final time points by deep sequencing.

An initial QC analysis of the screen results revealed several encouraging findings. First, hierarchical clustering of all of the sequenced samples—plasmid pools, T_0_ for each of the three cell lines, and T_f_ for each of the three cell lines—produced the patterns of clustering we would expect. Two replicates of the plasmid pool and the T_0_ sample for each of the three cell lines, for example, clustered closely together, suggesting that our sgRNA library was sufficiently represented across all three cell lines. All nine T_f_ samples also clustered appropriately, with the three parental samples gathering in one sub-cluster and the six samples belonging to the GR4 and WZR12 resistant derivatives gathering together in a separate sub-cluster (Supplementary Fig. 2a). These findings suggested to us that parental and drug-resistant cells respond differently to perturbations in the MCB-613 interactome, and that our screen was sufficiently designed and executed to detect these meaningful differences. Moreover, these findings were also recapitulated by a separate principal component analysis (PCA) (Supplementary Fig. 2b-c).

Moving forward, we reasoned that the target of MCB-613 would most likely be found among the genes which, when lost, conferred a greater growth disadvantage upon the resistant derivatives than upon the parental cells. Of all of these genes, it was the loss of Kelch-like ECH associated protein 1 (KEAP1) that most closely phenocopied the effects of MCB-613 and scored as the greatest collateral sensitivity to drug resistance in the screen (Fig. 3a, Supplementary Fig. 2d). Indeed, follow-up validation in parental PC9 and drug-resistant PFR3, GR4, and WZR12 cells confirmed that the loss of KEAP1 is differentially essential in the setting of EGFR inhibitor resistance (Fig. 3b). We took this as strong evidence that KEAP1 is in fact the protein interactor responsible for the selective cell death effect of MCB-613 in the setting of EGFR inhibitor resistance in NSCLC. From there, we further reasoned that, if it is in fact true that MCB-613 works through the inhibition of KEAP1, then the genetic knockout of this gene from a bulk population of resistant derivatives should eradicate all cells sensitive to the loss of KEAP1 and leave behind a reservoir of cells which no longer demonstrate any sensitivity to this molecule. As expected, genetic knockout of *KEAP1* rescued the selective cell death effects of MCB-613 across each of the resistant derivatives tested (Fig. 3c). Together, these results indicate that *KEAP1* is differentially essential in the setting of EGFR inhibitor resistance, and heavily suggest that the product of this gene is the protein interactor responsible for the collateral sensitivity of these cells to MCB-613.

**Fig. 3.**
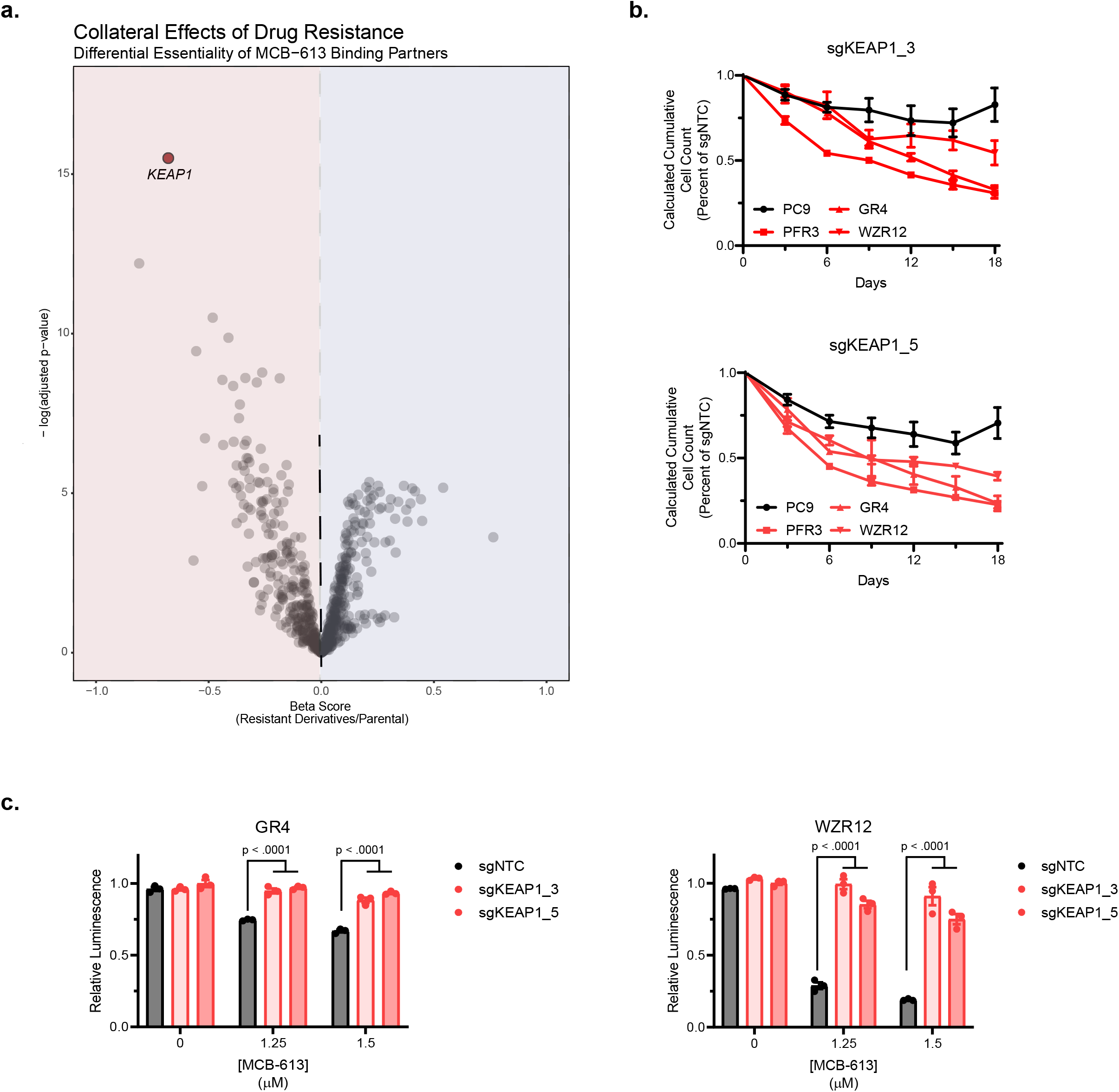
CRISPR/Cas9 loss-of-function screen and validation nominating KEAP1 as the target of MCB-613. a. Volcano plot depicting screen results. Each point represents relative essentiality of a gene encoding for a protein interactor of MCB-613 in drug-resistant (GR4 and WZR12) versus parental (PC9) cells. Negative and positive x-axis values indicate collateral sensitivities and resistances, respectively. b. Growth plots depicting calculated cumulative cell count over time in parental (PC9) and drug-resistant (PFR3, GR4, and WZR12) cells following knockout of *KEAP1* with sgKEAP1_3 (top) and sgKEAP1_5 (bottom). All results normalized to a nontargeting control. Data are mean ± SEM for n = 3 biologically independent experiments. c. Histograms depicting relative cell viability of drug-resistant GR4 (left) and WZR12 (right) cells after 72 hours of exposure to vehicle (DMSO) or two different doses of MCB-613 following knockout of *KEAP1* versus non-targeting control. Data are mean ± SEM for n = 3 biologically independent experiments.

### MCB-613 interacts directly with KEAP1

KEAP1 plays a canonical role as an adaptor subunit of the Cullin 3-based E3 ubiquitin ligase machinery. Under homeostatic conditions, KEAP1 homodimerizes and binds to the DLG and/or ETGE motifs of substrate proteins—most notably, the master transcription factor nuclear factor erythroid 2-related factor 2 (NRF2)—to facilitate their ubiquitination and subsequent degradation. In the presence of oxidative or electrophilic stress, however, the cysteine-rich KEAP1 senses these disturbances, dissociates from its substrate proteins, and allows them to accumulate. In the case of NRF2, this results in the activation of cell-protective, antioxidant transcriptional programs.^30^

We therefore hypothesized that MCB-613 interacts directly with KEAP1 to interrupt its degradation of substrate proteins, and that the accumulation of one of these substrates is responsible for the selective toxicity of this molecule in the setting of drug resistance. In order to test the first part of this hypothesis, we re-deployed ARM-3-124 in parental PC9 and drug-resistant WZR12 cells for one hour, before repeating the collection, cell lysis, and copper-catalyzed azide-alkyne cycloaddition steps which were described above. Rather than subjecting these samples to MS, however, we used an immunoblotting approach to observe whether the active derivative of MCB-613 pulled down KEAP1. As expected, ARM-3-124 bound to KEAP1 in both the parental cells and their drug-resistant derivatives, while the negative control ARM-3-115 showed little binding (Fig. 4a).

**Fig. 4.**
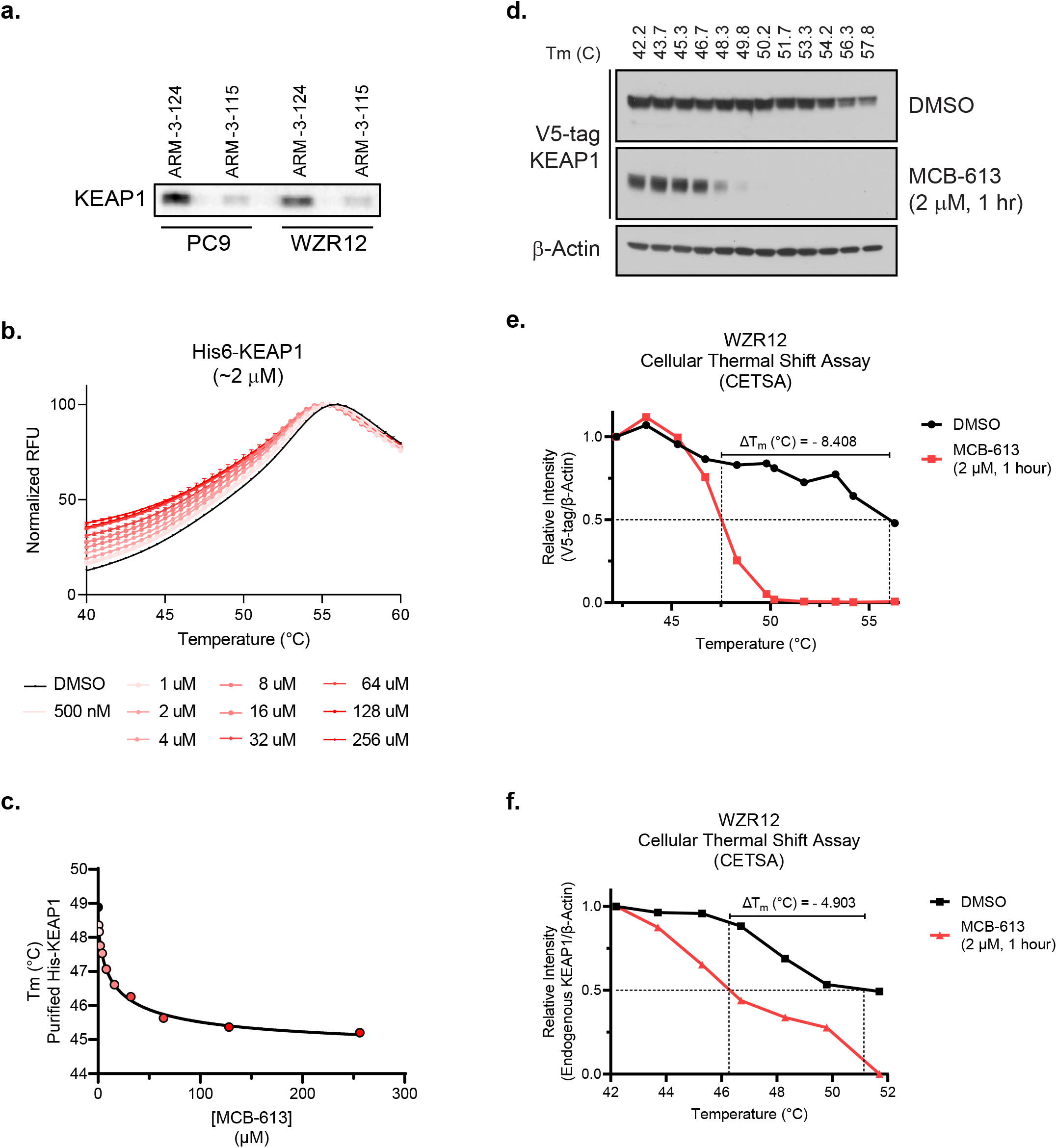
Evidence of a direct interaction between KEAP1 and MCB-613. a. Immunblot analysis of bound KEAP1 following treatment of parental (PC9) and drug-resistant (WZR12) cells with a 2 μM dose of either ARM-3-124 or ARM-3-115 for 1 hour followed by lysis, copper-catalyzed azide-alkyne cycloaddition, and gel electrophoresis. b. Thermal denaturation curve depicting fluorescence of SYPRO Orange induced by denaturation of purified His6-KEAP1 with increasing doses of MCB-613. Each condition internally normalized using non-linear fitting with a modified Boltzmann Equation. Data are mean ± SEM of n = 3 technical replicates. c. Dose-response curve depicting dose-dependent decrease in Tm of purified His6-KEAP1 with increasing doses of MCB-613. d. Immunoblot analysis of V5-tag following a cellular thermal shift assay in which lysates from drug-resistant WZR12 cells ectopically expressing V5-KEAP1 and treated with either vehicle (DMSO) or a 2 μM dose of MCB-613 for 1 hour were divided into aliquots and subjected to increasing temperatures to assess thermal stability of V5-KEAP1 as a proxy for direct binding. e. Line plot depicting densitometry analysis of immunoblot (d). f. Line plot depicting densitometry analysis of a similar CETSA study performed using endogenous KEAP1.

Next, we used several well-established techniques to confirm a direct interaction between MCB-613 and KEAP1. First, we used a pet28a-His6-KEAP1 construct to express wild-type human KEAP1 in *E. coli* before an Ni-NTA-based affinity purification.^31^ We then incubated purified His6-KEAP1 with increasing doses of MCB-613 in a SYPRO-Orange-based thermal shift assay.^32^ (In short, the thermal shift assay uses temperature gradients and hydrophobic dyes to detect melting temperature changes when purified proteins are incubated with suspect ligands.) As expected, this assay confirmed that MCB-613 interacts directly with purified His6-KEAP1; to our surprise, however, this resulted in an unusual, dose-dependent *decrease* in KEAP1 Tm (Fig. 4b-c). This suggests that, rather than stabilizing KEAP1 (in the manner of most small molecule ligands), MCB-613 decreases isothermal stability.

Although there is precedent for the destabilization of KEAP1 in thermal shift assays by small molecule ligands, we nevertheless decided to validate this somewhat counter-intuitive finding using orthogonal approaches.^33^ The cellular thermal shift assay (CETSA) operates according to the same basic premises as the traditional thermal shift assay, but in the setting of whole cell lysates and immunoblotting.^34^ We therefore expressed a V5-tagged KEAP1 in drug-resistant WZR12 cells and treated them with a physiological dose of MCB-613 for one hour before performing a CETSA. Consistent with our findings in the traditional thermal shift assay, CETSA confirmed that MCB-613 binds to KEAP1 and causes a *decrease* in melting temperature (Fig. 4d). We used densitometry analysis to calculate that MCB-613 reduced the Tm of V5-KEAP1 by more than 8.4 °C (Fig. 4e). To control for artifacts introduced by the use either of the V5 tag or the anti-V5 antibody, we recapitulated the CETSA and densitometry findings using an antibody that recognizes endogenous KEAP1 (Fig. 4f). Together, these data demonstrate the MCB-613 directly interacts with KEAP1, decreasing its thermal stability.

### MCB-613 promotes covalent dimerization of KEAP1

Because decreases in a protein’s stability often lead to degradation, we wondered whether MCB-613-mediated destabilization of KEAP1 led to a decrease in the latter’s protein levels. Thus, we briefly exposed parental PC9 and drug-resistant WZR12 cells to increasing doses of MCB-613 and probed for endogenous KEAP1. Although treatment with MCB-613 did in both cases induce a dose-dependent decrease in detectable KEAP1 levels by western blot, we also noticed the curious appearance of an additional band at roughly twice the estimated kilodalton molecular weight of monomeric KEAP1 (Fig. 5a).

**Fig. 5.**
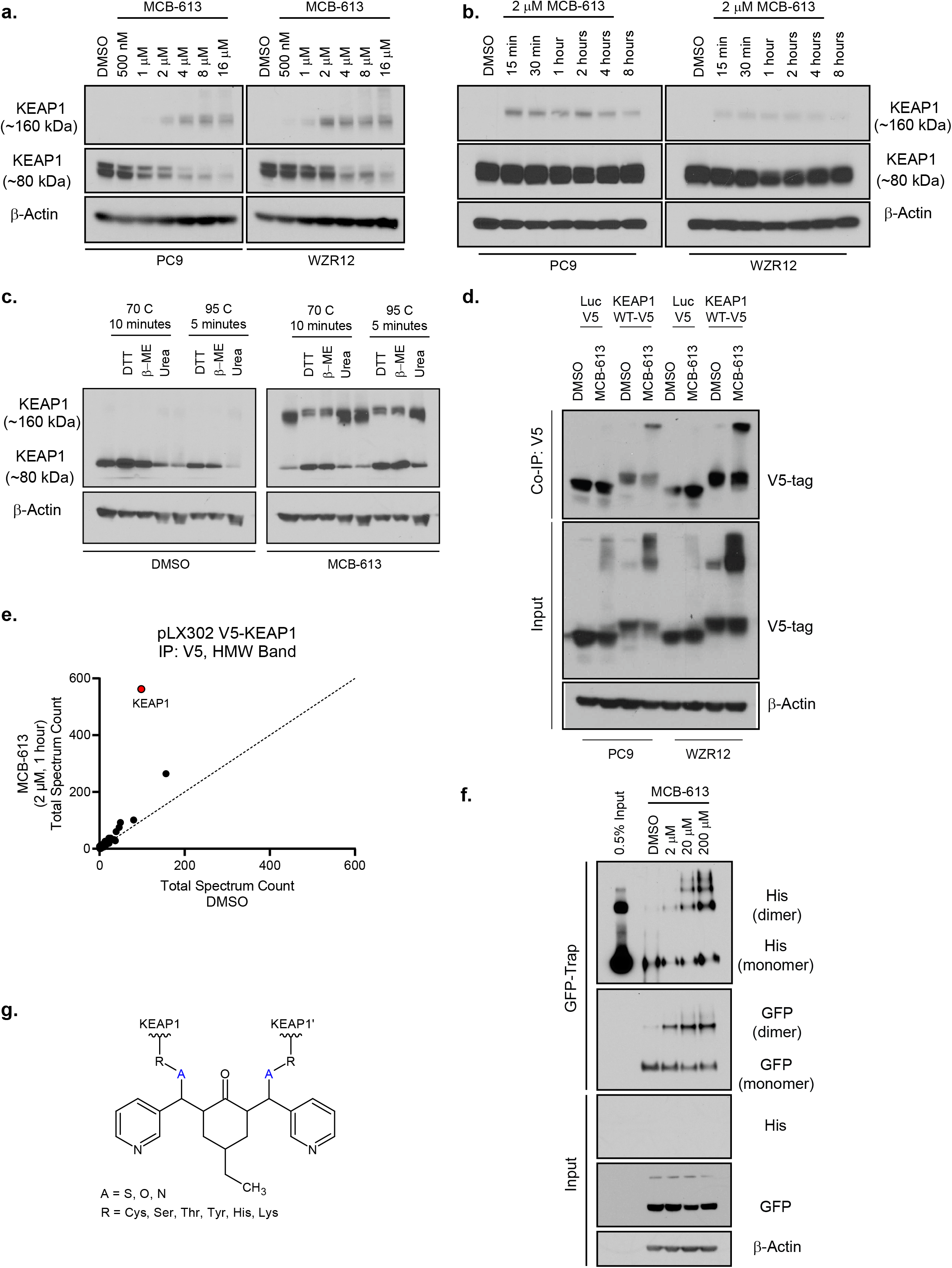
Evidence that MCB-613 induces covalent dimerization of KEAP1. a. Immunoblot analysis of endogenous KEAP1 levels at monomeric (~ 80 kDa) and dimeric (~ 160 kDa) molecular weights in parental (PC9) and drug-resistant (WZR12) cells after treatment with either vehicle (DMSO) or increasing doses of MCB-613 for 1 hour. b. Immunoblot analysis of endogenous KEAP1 levels at monomeric (~ 80 kDa) and dimeric (~ 160 kDa) molecular weights in parental (PC9) and drug-resistant (WZR12) cells over time after treatment with either vehicle (DMSO) or a 2 μM dose of MCB-613. c. Immunoblot analysis of endogenous KEAP1 levels at monomeric (~ 80 kDa) and dimeric (~ 160 kDa) molecular weights in drug-resistant WZR12 cells treated with either vehicle (DMSO) or a 2 μM dose of MCB-613 for 1 hour. Lysates were run under non-reducing versus reducing conditions and boiled at either 70 °C for 10 minutes or 95 °C for 5 minutes. d. Immunoblot analysis of V5 tag following immunoprecipitation of the V5 tag from parental (PC9) and drug-resistant (WZR12) cells ectopically expressing either a control V5-luciferase or V5-KEAP1 and treated with either vehicle (DMSO) or a 2 μM dose of MCB-613 for 1 hour. e. Scatterplot analysis of proteins identified by mass spectrometry to interact with V5-KEAP1. A high-molecular weight (~ 160 kDa), Coomassie-stained gel slice was obtained following immunoprecipitation of the V5 tag from drug-resistant WZR12 cells ectopically expressing V5-KEAP1 and treated with either vehicle (DMSO) or a 2 μM dose of MCB-613 for 1 hour. Slope of dotted line = 1 and represents hits equally identified between the conditions. f. Immunoblot analysis of Clover- and His-tagged KEAP1 after immobilization of HA-Clover KEAP1 on GFP-Trap beads and subsequent incubation with purified His6-KEAP1 and vehicle (DMSO) or increasing doses of MCB-613 for 15 minutes. g. Diagram depicting the proposed double Michael addition between KEAP1 and MCB-613.

Prior reports have shown that structurally diverse electrophiles known to interact covalently with KEAP1 (oltipraz, diethylmalate, sulforaphane, and tert-butylhydroquinone (TBHQ)) induce the conversion of KEAP1 from a primarily monomeric form, which migrates at ~ 68 kDa, to a largely dimeric or oligomeric form that can migrate at more than twice that molecular weight.^35^ We hypothesized that the loss of normal molecular weight KEAP1 that we observed in the immunoblotting experiment described above might be the result of a similar dimeric or even oligomeric transition to a higher molecular weight form of this protein. Indeed, further immunoblotting studies confirmed that treatment with a physiologically active dose of MCB-613 results not in the degradation of KEAP1, but in a rapid shift toward the dimeric molecular weight (Fig. 5b). Interestingly, these results persisted in the face of various reducing conditions, suggesting that this protein complex was not the result of traditional, non-covalent dimers or disulfide bridges (Fig. 5c).

To ensure that these results were not due to the artifactual appearance of some non-specific band, we confirmed these findings using ectopic V5-KEAP1 overexpression and co-immunoprecipitation with an anti-V5 antibody (Fig. 5d). Furthermore, excision, digestion, and mass spectrometry analysis of the high molecular weight band from the co-IP confirmed that it was substantially enriched for KEAP1 above all other protein species; this heavily suggested to us that the band represents a homo-, rather than hetero-, dimer (Fig. 5e). Lastly, an *in vitro* binding assay between two differentially tagged KEAP1 species treated with increasing doses of MCB-613 produced a similar transition from normal molecular weight to high molecular weight KEAP1 (Fig. 5f). Taken in their entirety, these results suggest that the appearance of this band represents the formation of a true, covalent homodimer of KEAP1. Because each molecule of MCB-613 bears not one but two electrophilic, α, β-unsaturated bonds, these results raise the intriguing possibility that a single molecule of MCB-613 may act as a double Michael acceptor to form a covalent bridge between two individual KEAP1 monomers (Fig. 5g).

### MCB-613 binds to KEAP1 within the dimerization domain to promote substrate accumulation

In order to better understand how MCB-613 induces covalent dimerization of KEAP1, we next used an HA-Clover tag system to design a variety of truncated forms of KEAP1. The first fragment, HA-Clover KEAP1^1-60^, is limited to the first sixty amino acids of KEAP1, which comprise the protein’s N-terminal domain. The second fragment, HA-Clover KEAP1^1-178^, further includes the protein’s BTB domain. Finally, the third and fourth fragments, HA-Clover KEAP1^1-314^ and HA-Clover KEAP1^1-597^, stretch to include the IVR and Kelch domains, respectively (Fig. 6a). We also included a full-length KEAP1 construct spanning the full 624 amino acids (HA-Clover KEAP1^WT^).

**Fig. 6.**
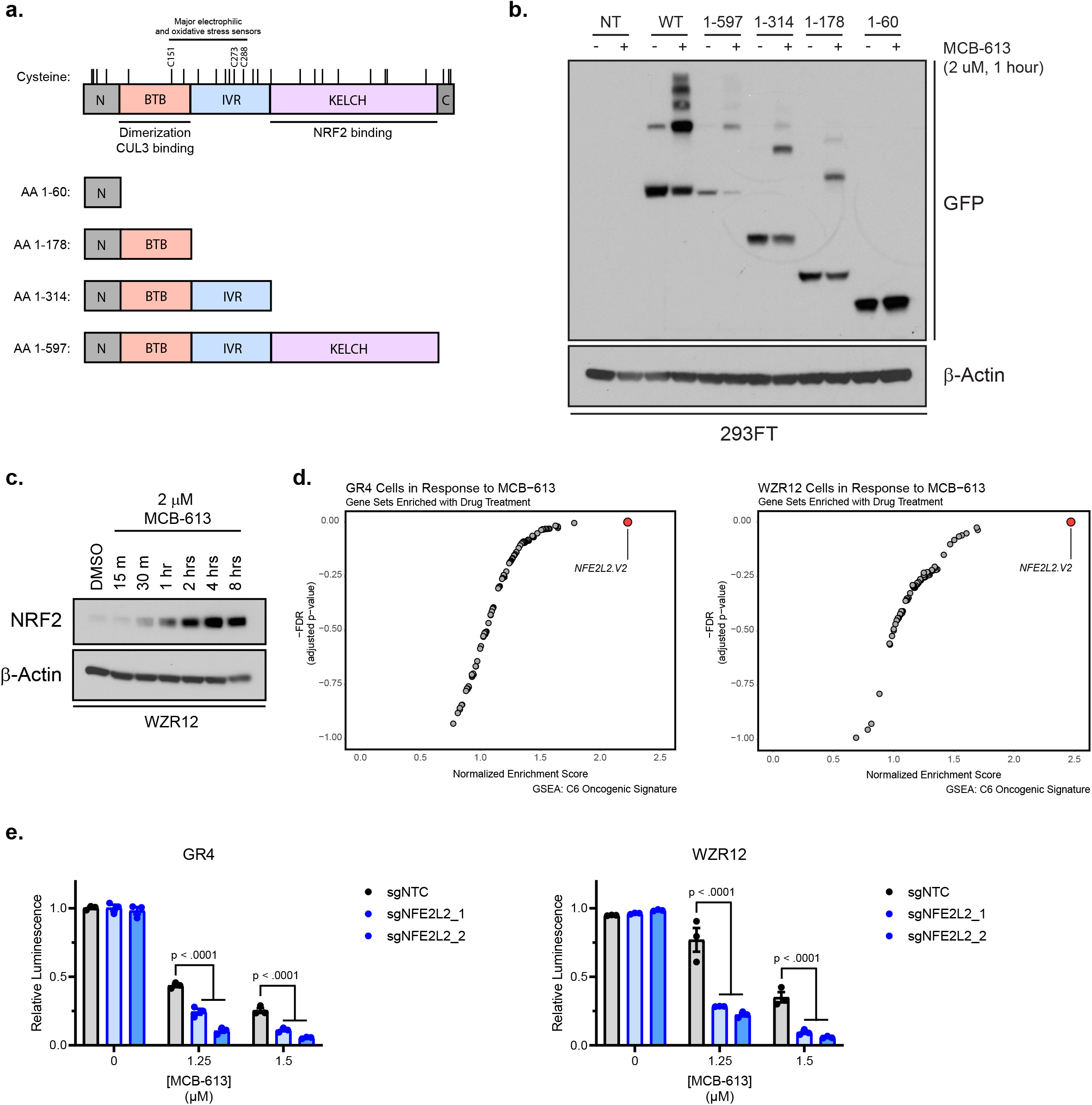
MCB-613 binds to KEAP1 within the dimerization domain to promote NRF2 accumulation and activation. a. Diagram depicting the protein domains of the full-length KEAP1 WT as well as the KEAP1^1-60^, KEAP1^1-178^, KEAP1^1-314^, and KEAP1^1-597^ truncation constructs. b. Immunoblot analysis of HA-Clover KEAP1 dimerization (upper bands) in 293FT cells transfected with (from left to right) no template, HA-Clover KEAP1^WT^, HA-Clover KEAP1^1-597^, HA-Clover KEAP1^1-314^, HA-Clover KEAP1^1-178^, and HA-Clover KEAP1^1-60^ and treated with vehicle (DMSO) or MCB-613 for one hour. c. Immunoblot analysis of NRF2 levels over time following treatment with a 2 μM dose of MCB-613 in drug-resistant WZR12 cells. d. Scatterplots depicting results of gene set enrichment analysis (GSEA) using the C6 Oncogenic Signature database in drug-resistant GR4 and WZR12 cells treated with a 2 μM dose of MCB-613 for 24 hours. d. Histograms depicting relative cell viability of drug-resistant GR4 (left) and WZR12 (right) cells after 72 hours of exposure to vehicle (DMSO) or two different doses of MCB-613 following knockout of *NFE2L2* versus non-targeting control. Data are mean ± SEM for n = 3 biologically independent experiments.

First, we transiently transfected each truncation construct into 293T cells. After 48 hours, we exposed each population to a physiological dose of MCB-613 for one hour, before collecting cells, lysing them, and immunoblotting with an anti-GFP antibody. As expected, MCB-613 caused the accumulation of a high molecular weight band with the HA-Clover KEAP1^WT^ construct. Interestingly, analogous bands appeared in the samples that were transfected with HA-Clover KEAP1^1-178^, HA-Clover KEAP1^1-314^, and HA-Clover KEAP1^1-597^ as well; indeed, the sole fragment which failed to reproduce this finding was HA-Clover KEAP1^1-60^ (Fig. 6b). This suggested to us that the covalent dimerization of KEAP1 induced by MCB-613 relies upon direct interaction with a residue located within the KEAP1 dimerization domain.

Given that KEAP1 relies upon homodimerization to function as a component of its E3 ubiquitin ligase complex, we wondered whether the covalent dimerization induced by MCB-613 affected protein function. We therefore treated drug-resistant WZR12 cells with a physiological dose of MCB-613 and observed that MCB-613-mediated KEAP1 inhibition indeed caused an accumulation of NRF2 over time (Fig. 6c). Furthermore, this NRF2 stabilization was associated with a strong upregulation of NRF2 transcriptional programs in both the drug-naïve and drug-resistant cell lines at 24 hours (Fig. 6d). Although NRF2 transcriptional programs are conventionally thought of as cytoprotective, recent reports have provided evidence that they can produce cytotoxicity in a subset of lung cancer cell lines through an NADH-mediated phenomenon known as “reductive stress”.^36^ Thus, we hypothesized that MCB-613-mediated KEAP1 inhibition might selectively target drugresistant NSCLC cells through the activation of NRF2 and initiation of reductive stress. To our surprise, however, a genetic knockout of NRF2 not only failed to rescue the collateral sensitivity of drug-resistant cells to MCB-613 but instead further sensitized them to it (Fig. 6e, Supplementary Fig. 3a). Although these results confirm, then, that MCB-613 inhibits the ability of KEAP1 to facilitate the degradation of target substrates such as NRF2, they also show that the NRF2 activation is dispensable for—and even counterproductive to—MCB-613-mediated cell death. This in turn provokes speculation that the collateral sensitivity of EGFR inhibitor-resistant cells to MCB-613-mediated inhibition of KEAP1 depends upon the accumulation of some alternative—and possibly heretofore undescribed—substrate of KEAP1.

## DISCUSSION

Here, we report that diverse mechanisms of drug resistance in *EGFR*-mutant nonsmall cell lung cancer converge on a collateral sensitivity to the understudied chemical compound MCB-613. Using a combination of unbiased, high-throughput –omics assays, we identify the molecular target of MCB-613 to be KEAP1. We also propose a compelling mechanism of action that involves covalent intermolecular bridging and is unrelated to the downstream activation of NRF2.

This work provides support for collateral sensitivity as an intellectual framework for approaching multifocal drug resistance, and nominates MCB-613 as a promising lead compound in the setting of *EGFR*-mutant NSCLC. Indeed, our data support a provocative notion: that cells of this genetically defined subtype can be forced into an “impossible decision” between sensitivity to EGFR inhibition on the one hand and sensitivity to MCB-613 on the other, and that, when faced with these two irreconcilable options, they can ultimately be overcome. As a matter of fact, several preliminary studies in other settings also speak to the exciting potential of this molecule as a promising lead compound: *in vivo* models of cardiac and nervous ischemia, for instance, reveal that MCB-613 is well tolerated and, thus far, free from obvious toxicities.^37,38^ In further support of our findings, moreover, these studies both also report that the use of MCB-613 leads to activation of NRF2 transcriptional programs. Although this was initially attributed to the downstream effects of steroid receptor coactivator (SRC) hyperstimulation, it is not lost on us that our reported mechanism of KEAP1 inhibition might also contribute to this phenotype.

Nevertheless, several key questions remain. First, the exact nature of the direct interaction between MCB-613 and KEAP1 remains opaque. Precise drug-target engagement dynamics are important not only for rational optimization of lead compounds such as this one, but also for prediction of MCB-613-specific drug resistance mechanisms which will inevitably surface. Second, it remains unclear which heretofore undescribed degradation substrate of KEAP1 accumulates and ultimately induces selective cell death in the setting of EGFR inhibitor resistance, and further investigation will be necessary to identify this factor and elucidate its mechanism. Finally, once such studies are complete, it will be important to establish the reasons for which EGFR inhibitor-resistant NSCLC cells become selectively sensitive to the modulation of this pathway.

This last point bears elaboration. It has long since been observed, for instance, that in the setting of non-small cell lung cancer, *EGFR* and *KEAP1* mutations are mutually exclusive.^39,40^ This remains poorly understood, and is further complicated by recent reports that genetic interruptions in the *KEAP1/NFE2L2/CUL3* axis may themselves contribute to the acquisition of EGFR inhibitor resistance.^41–43^ Even these fragments of understanding, however, are enough to indicate that a delicate and sensitive balance takes place between these two fundamental cellular pathways. Although our present findings now mean that this reciprocal relationship can be exploited at key times for therapeutic, anti-cancer gain, further investigation will be needed to more fully elucidate the exact nature of this relationship.

In all, our work nominates MCB-613 as an effective preliminary means of exploiting this precarious relationship, and provides an additional step toward understanding the complicated interplay at work here. More broadly, it also speaks to the power of collateral sensitivity as a framework for organizing studies which can both identify novel therapeutic strategies and provide insights into the nature of drug resistance itself.

## METHODS

### Cell lines and reagents

All cell lines were maintained in a humidified incubator at 37 °C with 5% CO_2_. 293FT cells were cultured in DMEM high glucose medium supplemented with 10% fetal bovine serum (FBS), 1% penicillin/streptomycin, 1% non-essential amino acids, 1% GlutaMAX, and 1% sodium pyruvate. All other cell lines were cultured in RPMI-1640 medium supplemented with 10% FBS and 1% penicillin/streptomycin. The PFR3, GR4, and WZR12 cell lines were a generous gift from Pasi Janne. The patient-derived MGH134 cell line was a generous gift from Aaron Hata. All other cell lines were purchased from the American Type Culture Collection (ATCC) or Duke Cell Culture Facility (CCF). Drug-resistant cell lines were derived as previously described.^9,44^ MCB-613 was a generous gift from Bert O’Malley. All other drugs were purchased from Cayman Chemical (gefitinib, osimertinib) and ChemDiv (C151-0598).

### High-throughput pharmacologic screen

PC9, PFR3, GR4, and WZR12 cells were seeded into drug-stamped, 384-well plates at a density of 300 cells per well. The ~2,100 compound library (Selleck Bioactives) was screened in duplicate at two different doses (2 and 10 μM) for each drug. After 72 hours, luminescence was read as a proxy for cell viability using Cell Titer Glo (Promega). The duplicate treatment wells for each compound were averaged and normalized to duplicate control wells from the same plate position. Normalized values were then processed by calculating the log2(resistant derivative/parental) for each compound at both doses. Hits which scored across multiple resistant derivatives and at the lower dose were prioritized.

### Dose-response (GI_50_) assays

Cells were seeded into 96-well plates at a density of 2,000 cells/well and allowed to adhere overnight. The following day, cells were treated in triplicate with either vehicle (DMSO) or an eight-point drug dilution and returned to the incubator. After 72 hours, luminescence was quantified with Cell Titer Glo (Promega). Relative cell viability was estimated by normalizing the raw luminescence values for the drug-treated wells to the DMSO controls.

### *In vivo* studies

Animal protocols were approved by the Animal Care and Use Committee of the Baylor College of Medicine. 5 x 10^6^ PC9 or GR4 cells were bilaterally implanted via subcutaneous injection into the flanks of 5-6 week-old female SCID mice and allowed to grow for two weeks. After two weeks, tumors were measured, mice were weighed, and treatment was initiated. Mice were treated with either saline or MCB-613 (20 mg/kg) in saline three times weekly via intraperitoneal (i.p.) injection. During the treatment period, mice were weighed and their tumors were measured weekly.

### Synthesis of ARM-3-124 and ARM-3-115

The preparation of chemical probes corresponding to C151-0598 and MCB-613 was developed in a two-step process. First, the methylene unit at the 4-position of the cyclohexanone was replaced with an oxygen atom (yielding intermediates ARM-3-126 and ARM-3-127) to ensure that this slight chemical modification did not alter the activity of the parent compounds (Supplementary Methods). Next, clickable activity-based probes harboring a propargyl handle at the 4-position of the cyclohexanone ring (ARM-3-115 and ARM-3-126) were prepared.

Briefly, the preparation of the target non-clickable (ARM-3-126, ARM-3-125) and clickable analogs (ARM-3-115, ARM-3-124) of C151-0598 and MCB-613 was performed in a one-step synthesis starting from either 4-methoxycyclohexanone or 4-propargylcyclohexanone. Using these starting materials, this step consisted in a double aldol condensation with benzaldehyde. It was achieved in ethanol at room temperature using potassium hydroxide as the base. Thereby, the non-clickable, ARM-3-126 and clickable, ARM-3-115, analogs of C151-0598 were obtained in modest to satisfactory yields (89 and 21%, respectively). The access to the analogs of MCB-613 were also obtained through a double aldol condensation of 1 and 2 with 3-picolinadehyde but under acidic conditions (HCl in acetic acid). The target compounds ARM-3-127 and ARM-3-124 were obtained in 71 and 37% yields, respectively.

For more detailed characterization of chemical synthesis, yield, and purity, please see Supplementary Methods.

### Copper-catalyzed azide-alkyne “click” reaction

The click chemistry experiments were conducted as previously described.^45^ Briefly, cells were seeded and exposed for one hour to a 2 μM dose of either the active ARM-3-124 or inactive ARM-3-115. After treatment, cells were washed three times in PBS and the pellet was resuspended in 1330 μL of 1 % NP40 buffer completed with protease inhibitor. Resuspended cells were sonicated for 30s three times and incubated 30 min on ice before centrifugation at 20,000 x g for 10 min. After cell lysis, proteins were extracted and quantified using the Bradford method. For each condition, up to 10 mg of protein in 9.4 ml 1 % NP40 buffer was used to perform the biotin addition using 100 μL of biotin azo-azide at 5 mM, 200 μL TCEP at 50 mM, 100 μL TBTA at 10 mM and 200 μL CuSO4 at 50 mM. Protein was then incubated for one hour in the dark at room temperature. After incubation, 40 ml ice-cold methanol was added. Protein was precipitated overnight in methanol at −20°C. Protein was then centrifuged at 5,200 x g for 30 min at 0°C and washed 3 times in the same conditions using ice-cold methanol. After drying, protein was resuspended in resuspension buffer (6 M urea, 2 M thiourea, 10 mM HEPES) and sonicated. Protein was supplemented with 40 μL DTT and incubated for 40 min at room temperature. 40 μL of iodoacetamide was added to the samples and incubated 30 min at room temperature in the dark. After pre-washing of the streptavidin beads, resuspended proteins were added to the beads in a 15 ml tube and incubated on a rotator for two hours at room temperature. Beads were collected by centrifugation at 200 x g for 3 min at room temperature and then washed with resuspension buffer (PBS and 1 % SDS, PBS) solution twice. After washing, elution was performed using sodium dithionite solution. Eluted proteins were incubated overnight at −20°C with ice-cold methanol. Proteins were then pelleted and dried before being resuspended in 4 % SDS buffer and 2X SDS free loading buffer. Samples were then separated by one-dimensional electrophoresis and stained with InstantBlue® dye. Gels were divided into pieces measuring approximately 1 mm^3^ and sent to Ross Tomaino at the Taplin Biological Mass Spectrometry Facility (Harvard Medical School) for identification of target proteins.

### Design of custom sgRNA library

The custom sgRNA library was designed using the list of 478 MS-identified genes. For each of these genes, 4 sgRNA sequences were taken from each of two previously published, full-genome CRISPR libraries.^46,47^ (Because sequences for a given gene were sometimes redundant between the two libraries, this approach produced an average of 7 unique sgRNA sequences per gene). Also included were 100 *AAVS1*-targeting “safe harbor” guides taken from a separate CRISPR library.^29^ Each sequence was appended with the same universal prefix and suffix sequences and synthesized commercially in pooled format (GenScript, Supplementary Table 1).

### Cloning of the CRISPR library

The commercially synthesized oligo pool was diluted 1:10 in water and amplified using the Phusion HotStart Flex polymerase (NEB) according to the manufacturer’s protocol with the Array_F and Array_R primers (Supplementary Table 1). The lentiCRISPR v2 vector (Addgene plasmid #52961) was digested with FastDigest Esp3I (Thermo Fisher) at 37 °C for 12 hours. The digested product was size selected with gel electrophoresis and extracted using the Zymo Gel DNA Extraction Kit. The eluted product was cleaned using the AMPure XP reagent at a 1:1 input-to-reagent ratio, and concentration was measured using the Qubit dsDNA HS Assay Kit (Invitrogen). The digested vector was combined with the amplified inserts in a 5:2 ratio by mass and ligated using a 2X Gibson Assembly Master Mix (NEB) at 50 °C for 30 minutes. The ligation product was cleaned using the AMPure XP reagent at a 1:1.4 input-to-reagent ratio and concentration was again measured using the Qubit dsDNA HS Assay Kit. The final product was transformed into electrocompetent cells (Lucigen), incubated overnight in LB broth at 37 °C, and isolated using the Qiagen Maxi Prep kit.

### Lentivirus production

Lentivirus for both the pooled CRISPR screen and individual sgRNAs was produced using 293FT cells. For every 1.3 μg of plasmid, 1 μg of the psPAX2 packaging vector and 0.657 μg of the VSVg envelope were diluted into 385 μl of Opti-MEM supplemented with 11.6 μl of the lipofectamine-2000 transfection reagent and 12.64 μl of PLUS reagent (Thermo Fisher). This mixture was added dropwise to 293FT cells, and cells were returned to the incubator for 4 hours. After the incubation, media was aspirated and replaced with an FBS-rich harvest medium (DMEM high glucose, 30% FBS, 1% PS, 1% NEAA, 1% GlutaMax, 1% sodium pyruvate) for 48 hours. Viruses were collected, filtered, and either used directly or aliquoted and stored at −80 °C. Viral titers were performed as previously described.^10^

### Pooled CRISPR screening

On Day 0, PC9, GR4, and WZR12 cells were seeded into 6-well plates at a density of 150,000 cells per well and allowed to adhere overnight. On Day 1, the cells were exposed to media containing library virus and the polybrene transduction reagent (8 μg/ml) before spinfection via centrifugation at 900 x g for 1 hour at room temperature. On Day 2, the media containing library virus and polybrene was removed and replaced with fresh media. On Day 3, the cells were subjected to selection with puromycin (2 μg/ml) for 48 hours. Then, on Day 5, the cells were resuspended, counted, divided into triplicates, and carried forward with passage every five days for an estimated 14 population doublings. Throughout the entirety of the screen, each replicate was represented at any given point in time by at least 4 x 10^6^ cells to guarantee at least 1000x coverage of the custom library. Pellets of at least 4 x 10^6^ cells were taken at T_0_, T_f_, and at each passage as well. Upon completion of the screen, pellets were lysed and genomic DNA was extracted using the QIAamp Blood Maxi Kit (Qiagen).

### CRISPR Screen sequencing and analysis

Extracted DNA was amplified with a two-step protocol. First, sgRNA libraries were PCR-amplified from genomic DNA using NEBNext Ultra II Q5 Master Mix according to the manufacturer’s protocol using the PCR 1 Forward and Reverse primers (Supplementary Table 1). Following the initial amplification, a second PCR step was undertaken to append the PCR 1 product for each condition with a unique combination of staggered forward and barcoded reverse primers to facilitate pooled sequencing (Supplementary Table 1, Supplementary Table 2). In all cases, the amplified libraries were purified with SPRIselect beads (Beckman Coulter) using right-sided selection. Samples were quantified with the Quant-iT dsDNA Broad Range Assay Kit (Thermo Fisher), pooled, and sequenced on an Illumina NextSeq 500 with 75 bp single-end sequencing. Analysis was performed using the Models-based Analysis of Genome-wide CRISPR/Cas9 Knockout (MAGeCK) software analysis pipeline under the default conditions. Data from the GR4 and WZR12 cell lines was collapsed into one condition representing drug resistance, and compared with data from the PC9 parental cell line.

### CRISPR screen validation by single gene knockout

To validate screening hits, the top two scoring sgRNA sequences for a given gene were taken from the custom library, cloned into the LCv2 vector, transfected into 293FT cells, and transduced into parental and drug-resistant NSCLC cells as described above (Supplementary Table 1). After 48 hours of puromycin selection, 350,000 cells per condition were seeded in triplicate into 10-cm dishes and passaged and counted every three days. Total cumulative cell counts over time were estimated using cell counts and doubling time while assuming exponential growth and a negligible death rate. Cumulative counts for a given sgRNA were normalized to cumulative counts for a non-targeting control.

### Protein purification

His6-KEAP1 was purified as previously described.^31^ Briefly, One Shot BL21 (DE3) chemically competent *E. coli* cells were transformed with the pET28a-His6-KEAP1 vector (Addgene plasmid #62454). Colonies were grown in LB broth containing 50 μM ZnCl_2_ and 50 μg/ml kanamycin at 37 °C until cultures reached an OD_600_ of 0.6-0.8 and then were allowed to cool to room temperature. Protein expression was induced with 250 μM IPTG and allowed to shake at 19 °C for 14 hours. Bacteria were collected and His6-KEAP1 purified on Ni-NTA agarose beads (Thermo Fisher) using the following buffers: lysis (50 mM NaH_2_PO_4_ (pH 8.0), 10 mM Imidazole, 5 mM β-mercaptoethanol (BME), 0.01% Triton X-100), washing buffer (50 mM NaH_2_PO_4_ (pH 8.0), 50 mM Imidazole, 500 mM NaCl, 5 mM BME, 0.01% Triton X-100), and elution (50 mM NaH_2_PO_4_ (pH 8.0), 125 mM Imidazole, 150 mM NaCl, 5 mM BME). Following elution, elution fractions were analyzed by gel electrophoresis to select and pool fractions of highest purity. The pooled fractions were then exchanged into storage buffer (50 mM Tris (pH 8.0), 10 mM DTT, 100 mM NaCl, 5% glycerol) using the Amicon Ultra-15 centrifugal filter column with a 10 kDa pore size (Millipore). Protein was quantified with the Bradford method before adjusting to a stock concentration of 4 mg/ml, aliquoting, flash freezing in liquid nitrogen, and storing at −80 °C.

### Thermal shift assay

The thermal shift assay was performed in a 384-well format as previously described.^48^ Purified His6-KEAP1 was diluted to a final concentration of 133 μg/ml (1.9 μM) in assay buffer (50 mM Na3PO4, 50 mM NaCl, pH 6.0) containing 8x SYPRO Orange diluted from a 5,000x stock (Invitrogen). The mixture was transferred to the wells of a 384-well plate and treated with either vehicle (DMSO) or doses of MCB-613 diluted in assay buffer. All conditions were plated in triplicate. The plate was sealed and subjected to centrifugation at 800 x g for 2 minutes to settle the samples before incubating at room temperature for 15 minutes. After the incubation ended, the plate was transferred to a Bio-Rad CFX384 Touch Real-Time PCR Detection System at 25 °C. The well temperature was increased in 0.5 °C increments with a 10-second equilibration time and fluorescence was measured by the built-in FRET channel until the machine reached 95 °C. For each condition, the melting temperature was determined by adjusting the raw values according to a modified Boltzmann Equation and calculating the average of the three replicates.

### Cellular thermal shift assay (CETSA)

The cellular thermal shift assay was performed according to previous reports.^34^ First, cells were seeded and allowed to adhere overnight. On the day of the assay, cells were treated with either vehicle (DMSO) or a 2 μM dose of MCB-613 for one hour before trypsinization, room temperature PBS washes, and collection in room temperature PBS supplemented with a protease inhibitor (Roche). For each condition, approximately 1 x 10^6^ cells were then transferred to each of 12 PCR strip tubes and sealed. A gradient PCR protocol including the following temperatures (°C) was initiated: 42.2, 43.7, 45.3, 46.7, 48.3, 49.8, 50.2, 51.7, 53.3, 54.2, 56.3, and 57.8. Samples were held off of the machine until the target temperature for each well was reached, and then placed on the machine and incubated at the target temperatures for 3 minutes. Each of the samples was then incubated in the thermal cycler at 25 °C for an additional 3 minutes, before being allowed to rest for a final 3 minutes at room temperature and on the bench top. Then, the PCR strip tubes were transferred to pre-chilled racks and snap frozen in liquid nitrogen, before incubating until completely thawed (~ 3-4 minutes) in a room temperature water bath. The cycle was repeated 3 more times for a total of 4 freeze-thaw cycles to promote cell lysis. After the final freeze-thaw cycle, the total volume of each PCR strip tube (~ 100 μl) was transferred to a 1.5 mL Eppendorf tube and centrifuged at 17,000 x g for 40 minutes at 4 °C. Debris pellets were discarded, and 60 μl of supernatant were transferred to a new tube containing 20 μl of NuPAGE LDS Sample Buffer (4X) and 4 μl of β-mercaptoethanol. Samples were vortexed and allowed to incubate at room temperature for 30-60 minutes before western blotting according to the procedures described below. Targets were detected using primary antibodies anti-KEAP1 clone 144 (MABS514, Sigma) and anti-V5 (#13202, CST) for endogenous and ectopic KEAP1, respectively.

### Densitometry analysis

Densitometry analysis of x-ray films from immunoblotting studies was performed using Adobe Photoshop and ImageJ. Images were converted to grayscale in Photoshop, and the intensity of bands was measured in ImageJ. For each blot, measurements were taken using a constant size selection for regions of interest. Background measurements were also collected from the blank regions located just above these bands. Pixel density was inverted by subtracting the measured intensities of both the band and background regions from 255. Net values were determined by subtracting the inverted densities of the background regions from the inverted densities of their corresponding bands of interest. The net value for each band of interest (e.g., KEAP1) was then normalized to the net value of the loading control for that lane (e.g., β-Actin). Finally, these normalized net values were themselves normalized to the 42.2 °C band in the CETSA experiment and plotted as a function of temperature.

### Western blot

Cells were washed in ice-cold PBS, collected via scraping, and re-suspended in ice-cold 1X Cell Lysis Buffer (CST) supplemented with protease and phosphatase inhibitors (Pierce). Cell suspensions were vortexed thoroughly and rotated end-over-end at 4 °C for 30 minutes before centrifugation at 14,800 rpm at 4 °C for 20 minutes. The debris pellets were discarded and the supernatants transferred to fresh Eppendorf tubes before quantification using the Bradford method. Samples were diluted to equal concentrations using lysis buffer and prepared for electrophoresis with NuPAGE LDS Sample Buffer (4X) before boiling at either 70 °C for 10 minutes or 95 °C for 15 minutes. Prior to electrophoresis, samples were reduced with DTT at a final concentration of 50 mM except where otherwise noted. Membranes were probed with primary antibodies anti-KEAP1 clone 144 (MABS514, Sigma), anti-NRF2 (Ab137550, Abcam), and anti-β-Actin (#4970, CST), and secondary antibodies anti-rabbit IgG, HRP-linked (#7074, CST) and anti-rat IgG, HRP-linked (#7077, CST).

### Co-immunoprecipitation

For co-IP studies, 4-5 x 10^6^ cells were seeded into two 15-cm dishes per condition and allowed to adhere overnight. The next day, cells were treated with a 2 μM dose of MCB-613 for one hour before scraping into ice-cold PBS. Cells were resuspended in IP buffer containing 40 mM Tris HCl (pH 7.4), 150 mM NaCl, 20 mM EDTA, 1 mM DTT, and 0.5% NP-40 supplemented with phosphatase and protease inhibitors, thoroughly vortexed, and rotated end-over-end at 4 °C for an hour. Samples were then subjected to centrifugation at 14,800 rpm for 20 minutes at 4 °C before harvesting the supernatant and measuring protein concentration using the Bradford method. A small volume of lysate was reserved for input, and the rest (600 – 800 μg) was adjusted so that a constant amount was loaded onto antibody-bound sepharose beads (see below) per sample. Samples were rotated end-over-end at 4 °C overnight. The following morning, samples were washed 5 times in ice-cold IP buffer before boiling at 95 °C for 5 minutes in NuPAGE LDS Sample Buffer (4X). Lysates were collected off of the beads using a 50 μl syringe (Hamilton) and analyzed via gel electrophoresis according to the procedures described above.

To prepare beads, 50 μl of Recombinant Protein G – Sepharose beads (Invitrogen) per condition were washed 3 times in IP buffer. The beads were then resuspended in IP buffer, and anti-V5-tag (E9H8O) Mouse mAb (#80076) (CST) was added at a 1:50 dilution as instructed by the manufacturer. The bead-antibody combination was then rotated endover-end at 4 °C for at least 5 hours. After the incubation, the beads were washed 3 times in IP buffer and divided evenly among the experimental conditions.

### Generation of HA-Clover KEAP1 constructs

Full-length KEAP1 WT was amplified out of the pET28a-His6-KEAP1 vector using the Phusion Hot Start Flex 2x Master Mix (NEB) according to the manufacturer’s instructions and appended with EcoRV and NotI restriction sites (for primer sequences, see Supplementary Table 1). Amplified full-length KEAP1 WT and the HA-Clover construct (Addgene plasmid #163336) were digested with the EcoRV and NotI enzymes (NEB) at 37 °C for 1 hour before heat inactivation at 65 °C for 5 minutes. Digestion fragments were purified by gel electrophoresis, excised, and extracted with the QIAQuick gel extraction kit (Qiagen). The fragments were combined using T4 DNA ligase (NEB) according to the manufacturer’s instructions. To generate KEAP1 truncations, site-directed mutagenesis was performed to introduce nonsense mutations at positions 61, 179, 315, and 598. (For primers, see Supplementary Table 1). All constructs were validated by Sanger sequencing (Eton Sequencing).

### Transient transfection of HA-Clover KEAP1 constructs

293FT cells were transiently transfected with 3 μg of each of the HA-Clover KEAP1 constructs diluted into Opti-MEM (Gibco) and supplemented with lipofectamine 2000 transfection reagent (Invitrogen). After 24 hours, media was refreshed. After another 24 hours, cells were treated with vehicle (DMSO) or 2 μM MCB-613 for one hour before collection in ice-cold PBS and preparation for western blot.

### GFP-Trap assay

293FT cells were transfected with HA Clover KEAP1 WT as described above. After 48 hours, the cells were washed once in ice-cold PBS and collected in ice-cold RIPA buffer (Sigma). The cells were incubated on ice for 30 minutes with intermittent vortexing, and the resultant lysates subjected to centrifugation at 10,000 rpm for 10 minutes at 4 °C. A fraction of the supernatant was reserved for input and prepared for western blotting as described above. The remaining supernatant was exposed to GFP-Trap agarose beads (Proteintech) and rotated end-over-end at 4 °C overnight before washing 5 times with icecold RIPA buffer. The beads were then equilibrated in room-temperature binding buffer (25 mM Tris (pH 7.5), 100 mM NaCl, 0.1% NP-40, 1 mM DTT, 5% glycerol). 10 μg of purified His6-KEAP1 was added and the samples were rotated end-over-end at room temperature for 15 minutes. MCB-613 (2, 20, or 200 μM) was added and the samples were rotated for another 15 minutes. Samples were washed 3 times in room-temperature binding buffer, re-suspended in 80 μl of NuPAGE LDS Sample Buffer (4X), boiled at 95 °C for 5 minutes, and analyzed by western blot. Targets were detected using anti-GFP (#2555, CST) and anti-His (#2365, CST) primary antibodies.

## ACKNOWLEDGMENTS

We would like to thank Pasi Janne (Harvard Medical School) for the PFR3, GR4, and WZR12 cell lines. We also thank Aaron Hata (Harvard Medical School) for the patient-derived MGH134 cell line. We thank Bert O’Malley (Baylor College of Medicine) for MCB-613, and for his continued support and feedback on this project. We thank Ross Tomaino (Taplin Mass Spectrometry Facility, Harvard Medical School) and Erik Soderblom (Proteomics and Metabolics Core Facility, Duke University School of Medicine) for their input and assistance on proteomic studies. Finally, we would like to thank Kevin H. Lin and Justine C. Rutter for their invaluable advice and support along the way. This work was supported by the NIH (R01CA263593 to K.C.W., K00CA245732-04 to J.H., F99CA264162 to B.M., F30CA247323 to C.C.S., and F31CA195967 to P.S.W.), the DOD (W81XWH-21-1-0362 to K.C.W.), the National Science Foundation (DGE-1106401 to G.R.A.), the Duke Medical Scientist Training Program (T32GM007171 to C.F.B., C.C.S., and S.T.K.), the Duke Graduate School (to C.F.B.), the Triangle Center for Evolutionary Medicine (to C.F.B.), and the Amgen Scholars Program (to S.G.).

