## Supplementary figures and images for "MCB-613 exploits a collateral sensitivity in drug resistant *EGFR*-mutant non-small cell lung cancer through covalent inhibition of KEAP1"

### Supplementary Figures

#### Supplementary Figure 1

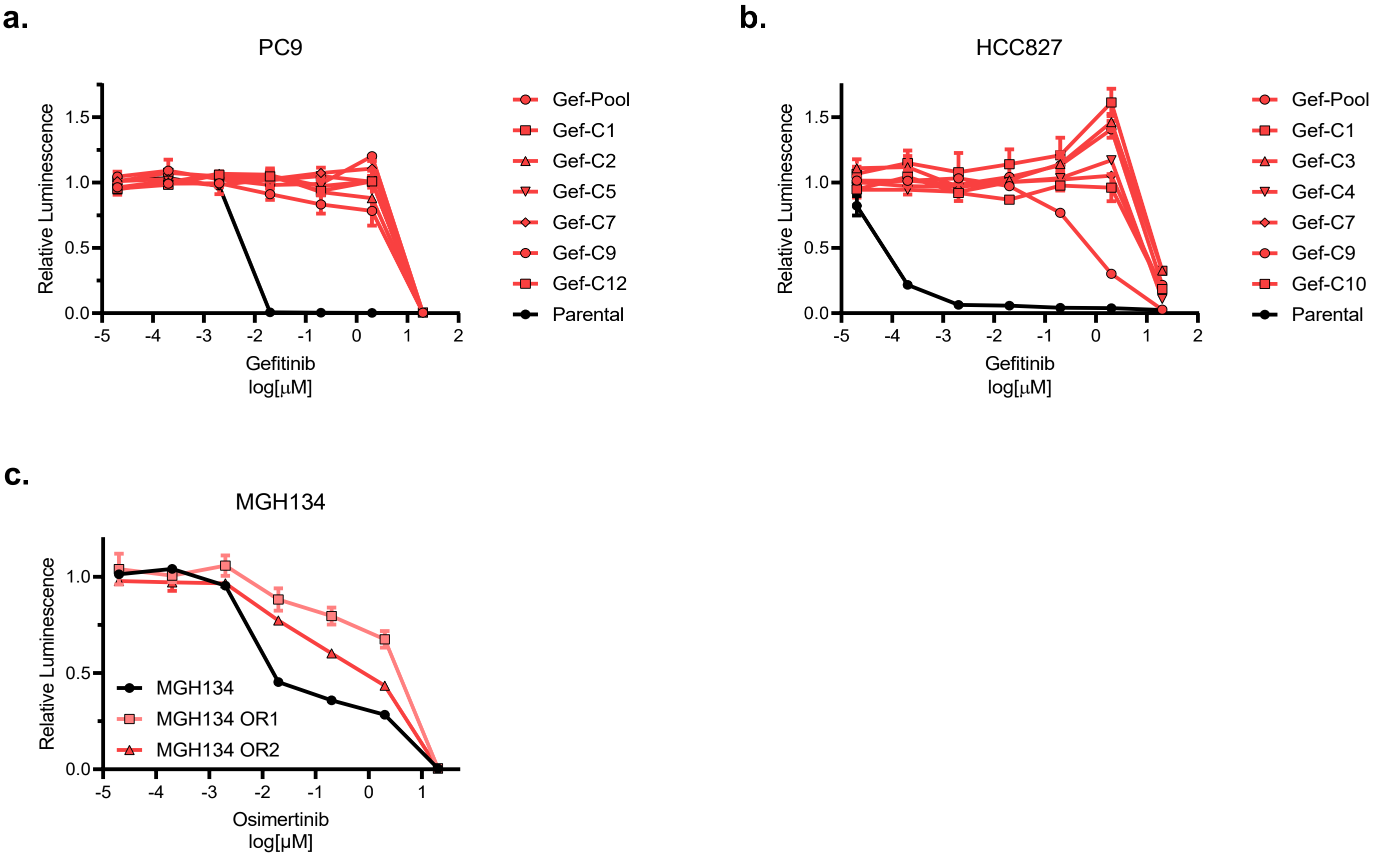

##### Supplementary Figure 2

**a.**

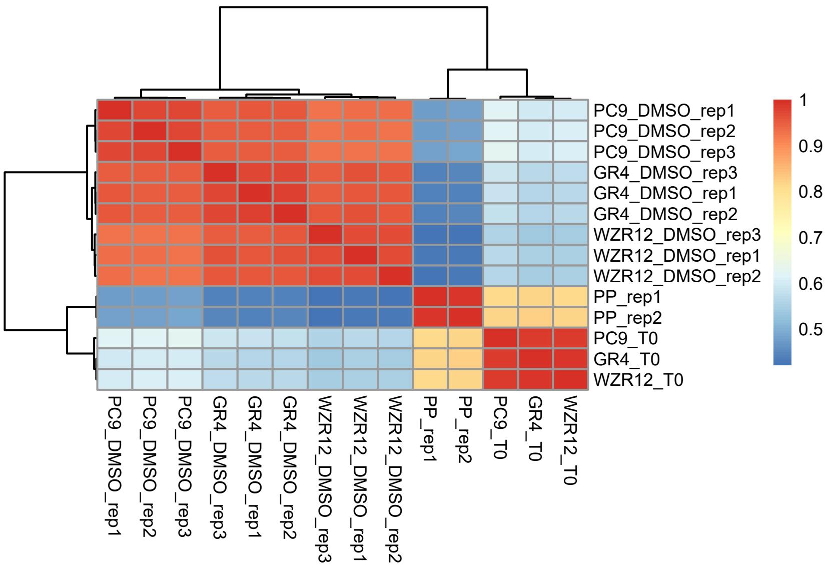


**b.**

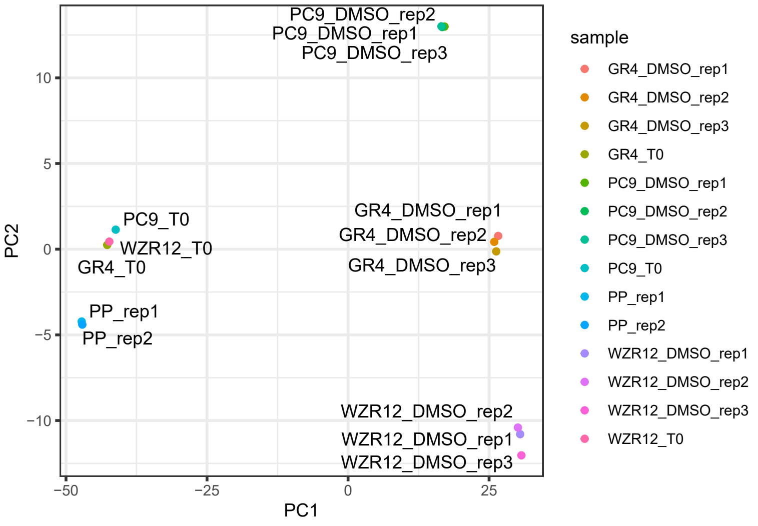


**c.**

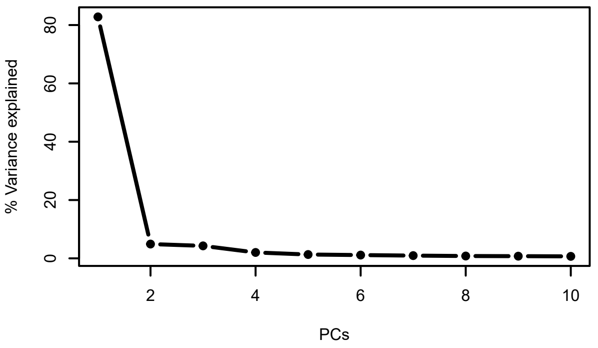


**d.**

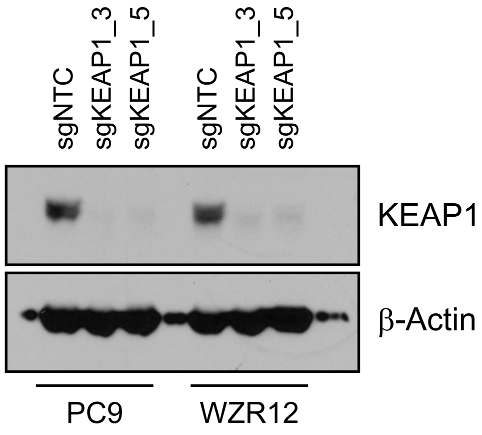

**Supplementary Figure 3**

**a.**

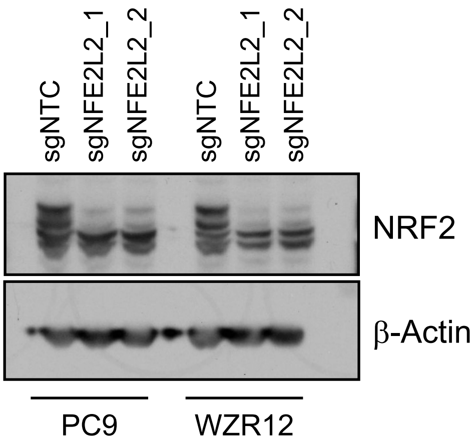
