## Supplementary Methods for "MCB-613 exploits a collateral sensitivity in drug resistant *EGFR*-mutant non-small cell lung cancer through covalent inhibition of KEAP1"

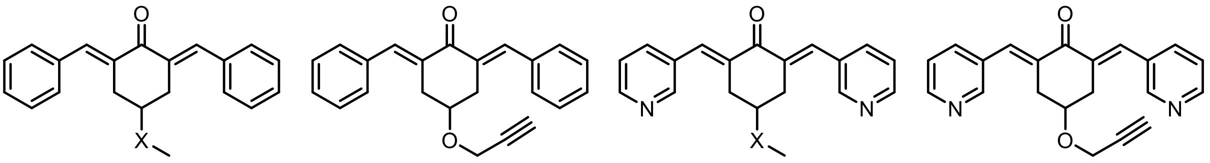

X = $CH_2$, **C151-0598**
X = O, **ARM-3-126**

**ARM-3-115**

X = $CH_2$, **MCB-613**
X = O, **ARM-3-127**

**ARM-3-124**

Structure of **C151-0598**, **MCB-613** and their respective clickable and non-clickable analogs.

The preparation of chemical probes corresponding to **C151-0598** and **MCB-613** was developed in a two-step process. First, the replacement of the methylene unit on the 4-position of the cyclohexanone with an oxygen atom (**ARM3-126** and **ARM-3-127**) was undertaken to ensure that this slight chemical modification did not alter the activity of the parent compounds. Thereafter, clickable activity-based probes harboring a propargyl handle on the 4-position of the cyclohexanone ring were prepared, **ARM-3-115** and **ARM-3-124**.

The preparation of the target non-clickable (**ARM-3-126**, **ARM-3-125**) and clickable analogs (**ARM-3-115**, **ARM-3-124**) of **C151-0598** and **MCB-613** was performed in a one-step synthesis starting from either 4-methoxycyclohexanone, **1** or 4-propargylcyclohexanone, **2**. Using these starting materials, this step consisted in a double aldol condensation with benzaldehyde. It was achieved in ethanol at room temperature using potassium hydroxide as the base. Thereby, the non-clickable, **ARM-3-126** and clickable, **ARM-3-115**, analogs of **C151-0598** were obtained in modest to satisfactory yields (89 and 21%, respectively). The access to the analogs of **MCB-613** were also obtained through a double aldol condensation of **1** and **2** with 3-picolinadehyde but under acidic conditions (HCl in acetic acid). The target compounds **ARM-3-127** and **ARM-3-124** were obtained in 71 and 37% yields, respectively.

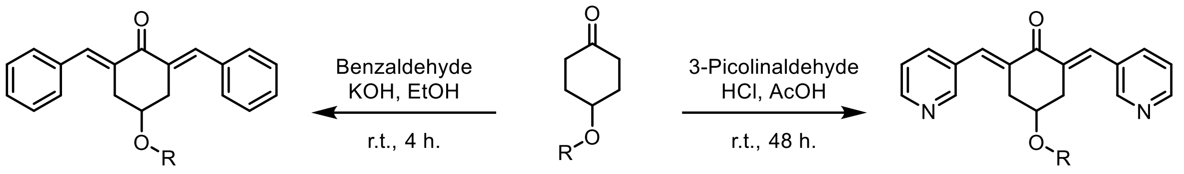


**ARM-3-126**, R = Me, 89%
**ARM-3-115**, R = $CH_2CCH$, 21%

**1**, R = Me
**2**, R = $CH_2CCH$

**ARM-3-127**, R = Me, 71%
**ARM-3-124**, R = $CH_2CCH$, 37%

Preparation of the clickable and non-clickable probes **ARM-3 -115, -124, -126** and **-127**.

General considerations:

All organic solvents were purchased from commercial sources and used as received. All chemicals were purchased from Merck or Acros and used without further purification; 4-(propargyloxy)cyclohexanone (compound **2**) was prepared according to Glaxosmithkline patent WO2007/107565, 2007 A1. Thin layer chromatography (TLC) was performed on precoated Merck 60 GF254 silica gel plates and revealed by spraying (*p*-anisaldehyde/$H_2SO_4$), and detection by means of UV light at 254 nm. $^1H$ and $^{13}C\{^1H\}$ NMR spectra were recorded on a Bruker Advance 400 MHz spectrometer; chemical shifts are given in ppm and referenced to the solvent residual peak ($CDCl_3$: 7.16 ppm for $^1H$ and 77.16 for $^{13}C\{^1H\}$); coupling constants are given in Hertz (Hz). The purity of the final compounds (**ARM-3-115, -124, -126** and **-127**) was verified to be ≥ 95 % purity by HPLC analysis at 254 nm on a JASCO PU-2089 equipped with a UV detector, Ascentis Express Suppelco column (5 µm, RP-C18) 4.6 mm × 100 mm; step-wise gradient water/acetonitrile (1‰ formic acid): 100/0 during 1 min then 100/0 to 0/100 in 5 min and then a plateau at 0/100 for 1 min before returning to initial conditions 100/0 over 1 min. (analysis duration: 8 min.); flow: 2 mL/min. High resolution mass spectra (HRMS) were recorded on ESI-LTQ Orbitrap U3000 RSLC spectrometer.

Preparation of 2,6-di((*E*)-benzylidene)-4-(prop-2-yn-1-yloxy)cyclohexanone (ARM-3-115):

*Synthesis*: To a solution of 4-(propargyloxy)cyclohexanone (compound **2**, 100 mg, 0.66 mmol, 1.0 equiv.), in ethanol (1.0 mL) at room temperature, were successively added benzaldehyde (139 mg, 1.31 mmol, 2.0 equiv.), and potassium hydroxide (73 mg, 1.31 mmol, 2.0 equiv.). The reaction mixture was stirred at room temperature until complete consumption of the starting materials (TLC monitoring, *ca.* 4 h). The reaction mixture was then concentrated to dryness and the crude material was purified over a silica gel column chromatography using a mixture of cyclohexane and ethyl acetate as the eluent. The title compound was obtained as a white solid in a 21% yield (45 mg).

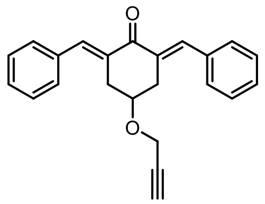

*Characterization*: $^1$H NMR ($CDCl_3$, 400 MHz) δ 7.89 (s, 2H), 7.48-7.34 (m, 10H), 4.06 (d, *J* = 2.2 Hz, 2H), 4.01 (ddd, *J* = 10.8, 7.3, 3.5 Hz, 1H), 3.21 (d, *J* = 15.0 Hz, 1H), 3.09 (dd, *J* = 16.4, 7.1 Hz, 1H), 2.30 (t, *J* = 2.0 Hz, 1H); $^{13}$C{$^1$H} NMR ($CDCl_3$, 101 MHz) δ 188.8, 139.2, 135.7, 132.3, 130.4, 129.0, 128.6, 79.6, 74.7, 71.6, 55.7, 33.5; HRMS (ESI+): calcd for $[M+H]^+$ $C_{23}H_{21}O_2$, 329.1536, found 329.1532; HPLC ($\lambda_{254}$) 97 % purity.

Preparation of 2,6-di((*E*)-benzylidene)-4-methoxycyclohexanone (ARM-3-126):

*Synthesis*: To a solution of 4-methoxycyclohexanone (compound **1**, 100 mg, 0.78 mmol, 1.0 equiv.), in ethanol (1.0 mL) at room temperature, were successively added benzaldehyde (166 mg, 1.56 mmol, 2.0 equiv.), and potassium hydroxide (88 mg, 1.56 mmol, 2.0 equiv.). The reaction mixture was stirred at room temperature until complete consumption of the starting materials (TLC monitoring, *ca.* 16 h). The reaction mixture was then filtered on a glass frit (porosity 4) and the solids were washed with a minimum (1 mL, twice) of ice-cold ethanol. The resulting white powder was dried under high vacuum to afford the title compound in an 89% yield (210 mg).

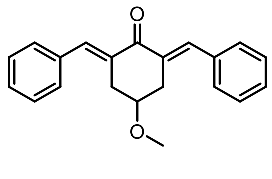

*Characterization*: $^{1}$H NMR ($CDCl_3$, 400 MHz) δ 7.89 (s, 2H), 7.48-7.33 (m, 10H), 3.64 (tt, *J* = 7.1, 3.4 Hz, 1H), 3.22 – 3.15 (m, 5H), 3.10 – 3.04 (m, 2H); $^{13}C\{^{1}H\}$ NMR ($CDCl_3$, 101 MHz) δ 189.1, 139.0, 135.8, 132.6, 130.4, 128.9, 128.6, 74.0, 56.2, 33.3; HRMS (ESI+): calcd for $[M+H]^+$ $C_{21}H_{21}O_2$, 305.1536, found 305.1533; HPLC ($\lambda_{254}$) 95 % purity.

Preparation of 4-(propargyloxy)-2,6-bis(3-picolilidene)cyclohexanone (ARM-3-124):

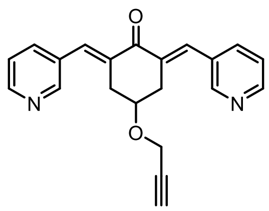


*Synthesis*: To a solution of 4-propargyloxycyclohexanone (compound **2**, 100 mg, 0.66 mmol, 1.0 equiv.), in acetic acid (1.5 mL) at room temperature, was added 3-picolinaldehyde (140 mg, 1.31 mmol, 2.0 equiv.), before pure hydrochloric acid (gas) was bubbled into the reaction media during 2 h. The reaction mixture was left to stir at room temperature until complete consumption of the starting materials (TLC monitoring, *ca.* 6 days). The reaction mixture was then diluted with water 20 mL and basified to pH 12 by adding a 2 M solution of sodium hydroxide. The media was extracted with ethyl acetate (50 mL, 3 times) and the combined organic layers were washed once with saturated aqueous $NaHCO_3$ and once with brine. The organic layer was dried over magnesium sulphate and the volatiles were evaporated off. The resulting crude material was eventually purified on a silica gel column chromatography using mixtures of ethyl acetate and cyclohexane as the eluent. Thereafter, the compound was recrystallized from a dichloromethane/ether solution and dried under high vacuum. The title compound was obtained as a colourless solid in a 37% yield (80 mg).

*Characterization*: $^{1}$H NMR ($CDCl_3$, 400 MHz) δ 8.73 (d, *J* = 1.8 Hz, 2H), 8.59 (dd, *J* = 4.8, 1.4 Hz, 2H), 7.84 (s, 2H), 7.79 – 7.77 (m, 2H), 7.38 (dd, *J* = 7.9, 4.9 Hz, 1H), 4.13 – 4.08 (m, 1H), 4.04 (d, *J* = 2.4 Hz, 2H), 3.21 – 3.10 (m, 2H), 2.30 (t, *J* = 2.4 Hz, 1H);

$^{13}C\{^1H\}$ NMR ($CDCl_3$, 101 MHz) δ 187.8, 151.2, 149.7, 137.1, 135.8, 134.0, 131.5, 123.5, 79.3, 75.0, 70.8, 55.8, 33.2; HRMS (ESI+): calcd for $[M+H]^+$ $C_{21}H_{19}O_2N_2$, 331.1441, found 331.1436; HPLC ($\lambda_{254}$) 95 % purity.

Preparation of 4-(propargyloxy)-2,6-bis(3-picolilidene)cyclohexanone (ARM-3-127):

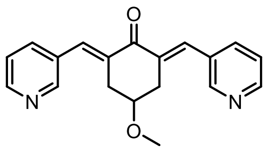


*Synthesis*: To a solution of 4-methoxycyclohexanone (compound **1**, 100 mg, 0.78 mmol, 1.0 equiv.), in acetic acid (1.0 mL) at room temperature, was added 3-picolinaldehyde (167 mg, 1.56 mmol, 2.0 equiv.), before pure hydrochloric acid (gas) was bubbled into the reaction media during 2 h. The reaction mixture was left to stir at room temperature until complete consumption of the starting materials (TLC monitoring, *ca.* 48 h.). It was then diluted with 1.0 mL of acetone and let to stand still for 2 h; the pure product slowly crystallises and can be collected by centrifugation and further washed with ether (twice, 2 mL). Thereafter, the solid product was solubilized in water (20 mL) and pH was adjusted to 12 using a 2 M solution of sodium hydroxide. Next, the product was extracted using ethyl acetate (20 mL, 3 times) and the combined organic layers were washed twice with saturated aqueous $Na_2CO_3$ and once with brine. The organic layer was dried over magnesium sulphate and the volatiles were evaporated off to afford the title compound as a colourless solid in a 71% yield (170 mg).

*Characterization*: $^1H$ NMR ($CDCl_3$, 400 MHz) 8.72 (d, $J$ = 1.7 Hz, 2H), 8.58 (dd, $J$ = 4.8, 1.3 Hz, 2H), 7.83 (s, 2H), 7.76 (d, $J$ = 7.9 Hz, 2H), 7.36 (dd, $J$ = 7.9, 4.9 Hz, 2H), 3.72 (tt, $J$ = 6.2, 3.3 Hz, 1H), 3.19 (s, 3H), 3.17 - 3.05 (m, 4H); $^{13}C\{^1H\}$ NMR ($CDCl_3$, 101 MHz) δ 188.1, 151.1, 149.6, 137.2, 135.6, 134.3, 131.6, 123.5, 73.1, 56.2, 33.0; HRMS (ESI+): calcd for $[M+H]^+$ $C_{19}H_{19}O_2N_2$, 307.1441, found 307.1436; HPLC ($\lambda_{254}$)

98 % purity.

**Copies of $^1$H & $^{13}$C{$^1$H} NMR and HRMS spectra and UV-HPLC traces: ARM-3-115**

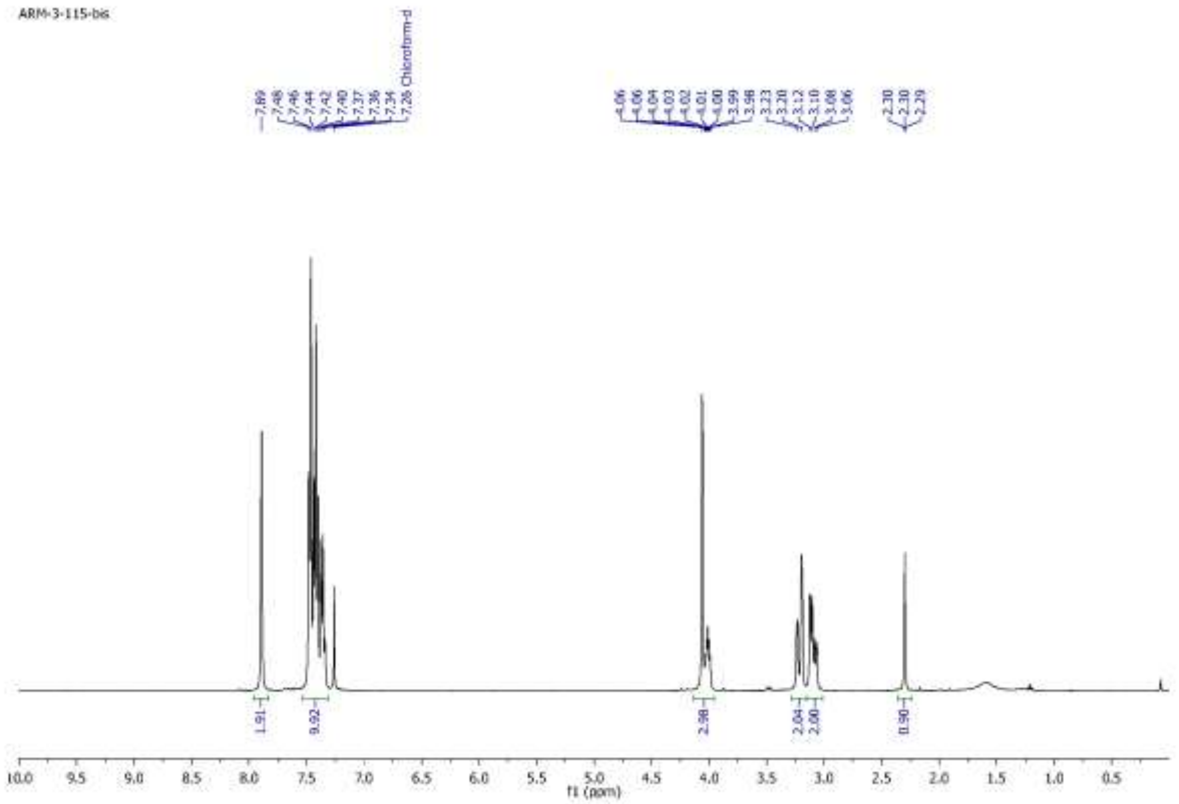


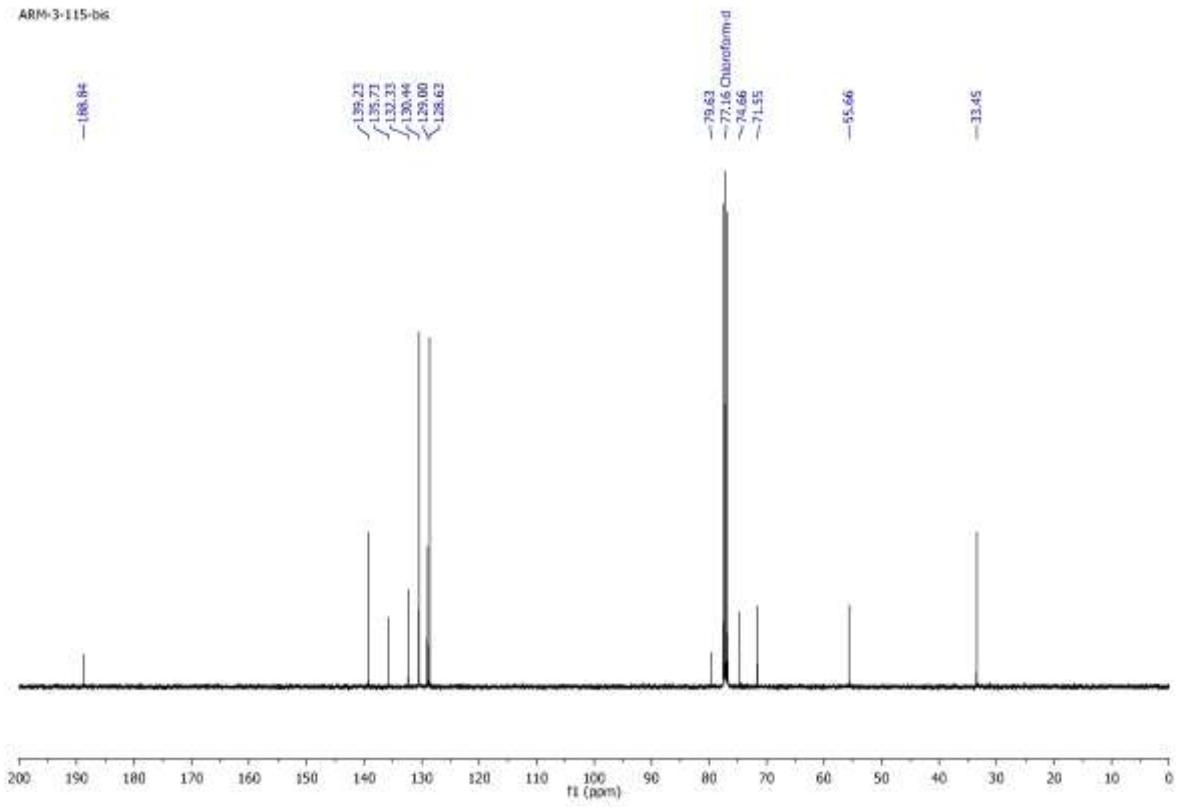

ARM3-115 #1557-2310 RT: 12.25-18.08 AV: 377 NL: 5.06E8
T: FTMS + p ESI Full ms [132.0000-1200.0000]

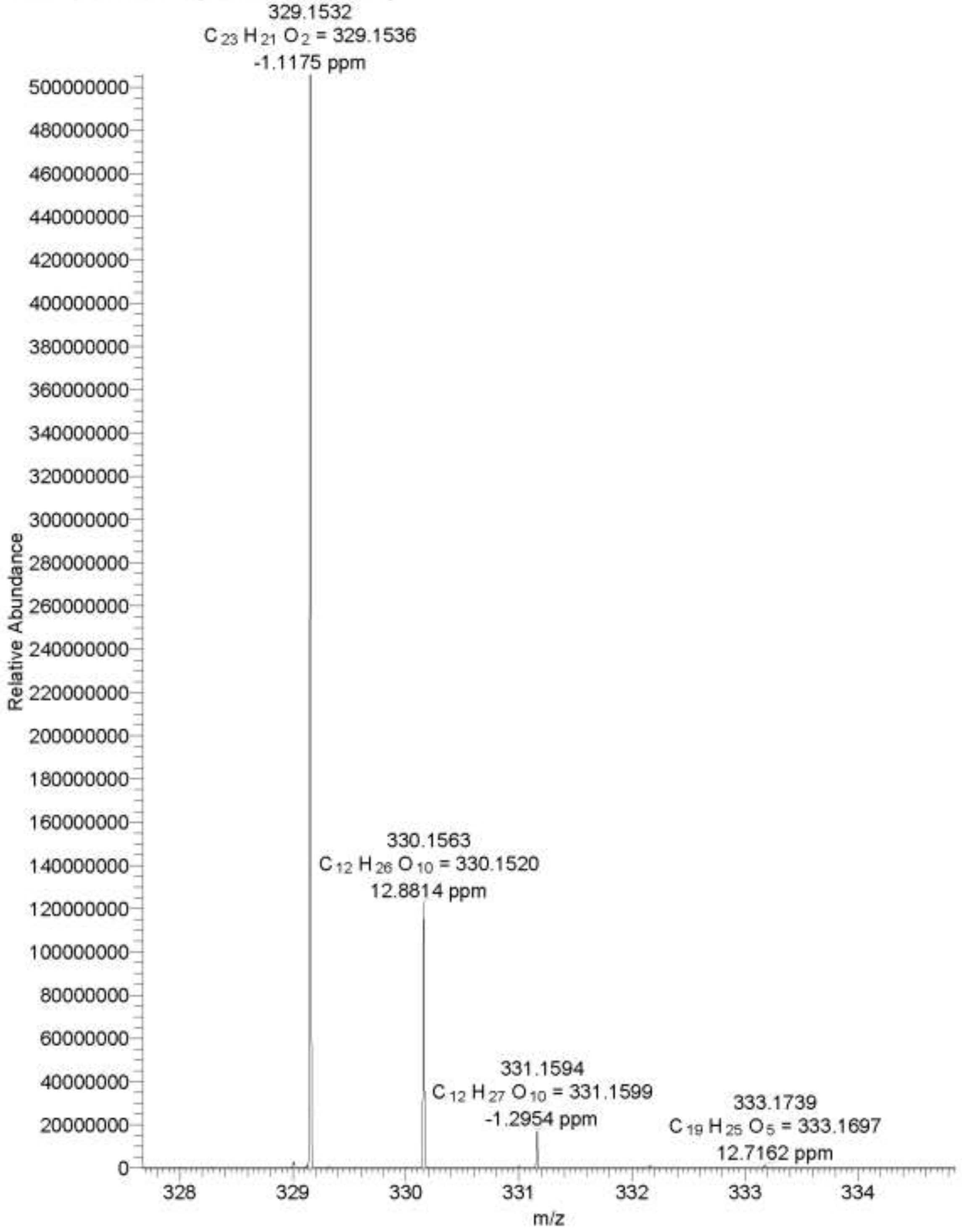

*ARM-3-115 ARM-3-115 29/07/2020 11:05:53*

Chromatogram

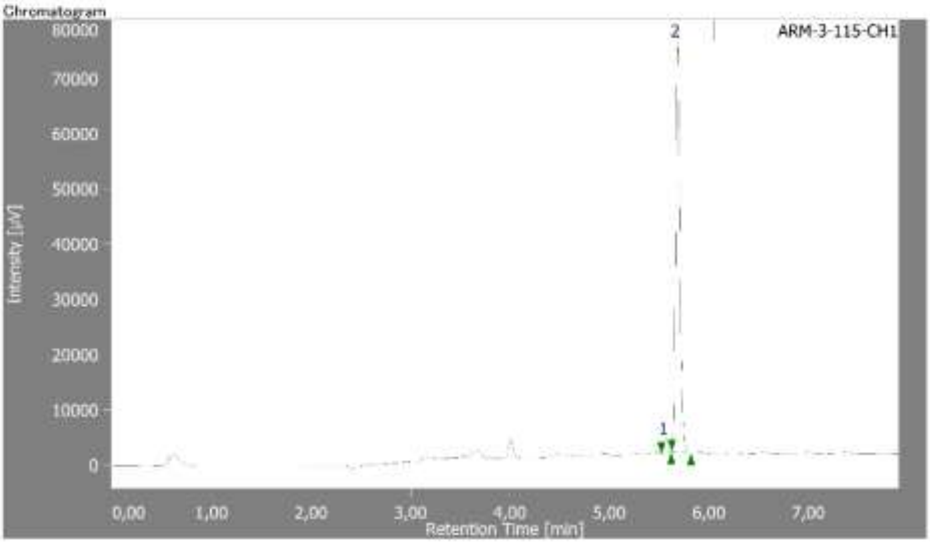


**Peak Information**

| # | Peak Name | CH | tR [min] | Area [µV·sec] | Height [µV] | Area% | Height% | Quantity | NTP | Resolution | Symmetry Factor | Warning |
|---|---|---|---|---|---|---|---|---|---|---|---|---|
| 1 | Unknown | 1 | 5.567 | 7336 | 2718 | 3.076 | 3.526 | N/A | 84833 | 1.494 | 1.107 | |
| 2 | Unknown | 1 | 5.683 | 231140 | 74362 | 96.924 | 96.474 | N/A | 80702 | N/A | 1.135 | |

Gradient Profiles - No data.

### ARM-3-126

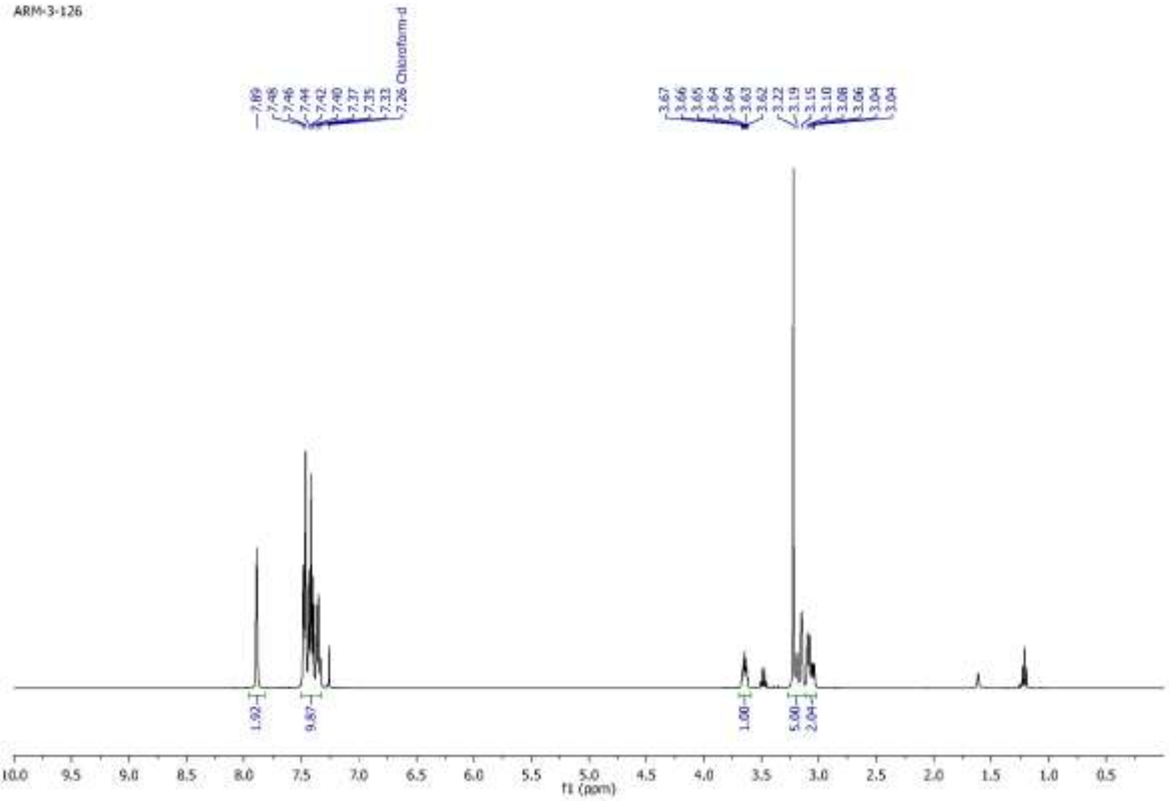
ARM-3-126
7.89
7.48
7.46
7.44
7.42
7.40
7.37
7.35
7.33
7.26 Chloroform-d
3.67
3.66
3.65
3.64
3.64
3.63
3.62
3.22
3.19
3.15
3.10
3.08
3.06
3.04
3.04
1.92
9.87
1.00
5.00
2.04
f1 (ppm)

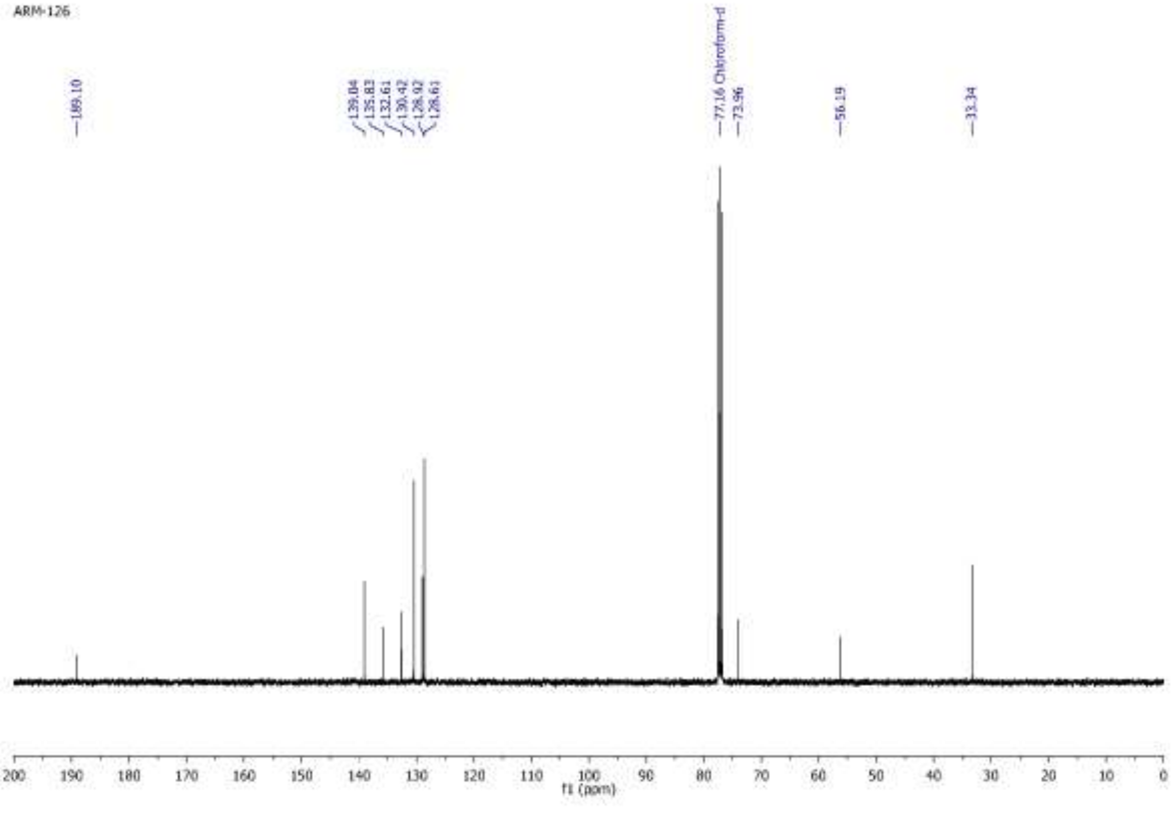
ARM-126
189.10
139.04
135.83
132.61
130.42
128.92
128.61
77.16 Chloroform-d
73.96
56.19
33.34
f1 (ppm)

C:\Users\...\Desktop\HRMS Mauro\ARM3-126 9/13/2020 9:35:05 AM

ARM3-126 #5-2792 RT: 0.04-22.00 AV: 1394 NL: 5.82E8
T: FTMS + p ESI Full ms [132.0000-1200.0000]

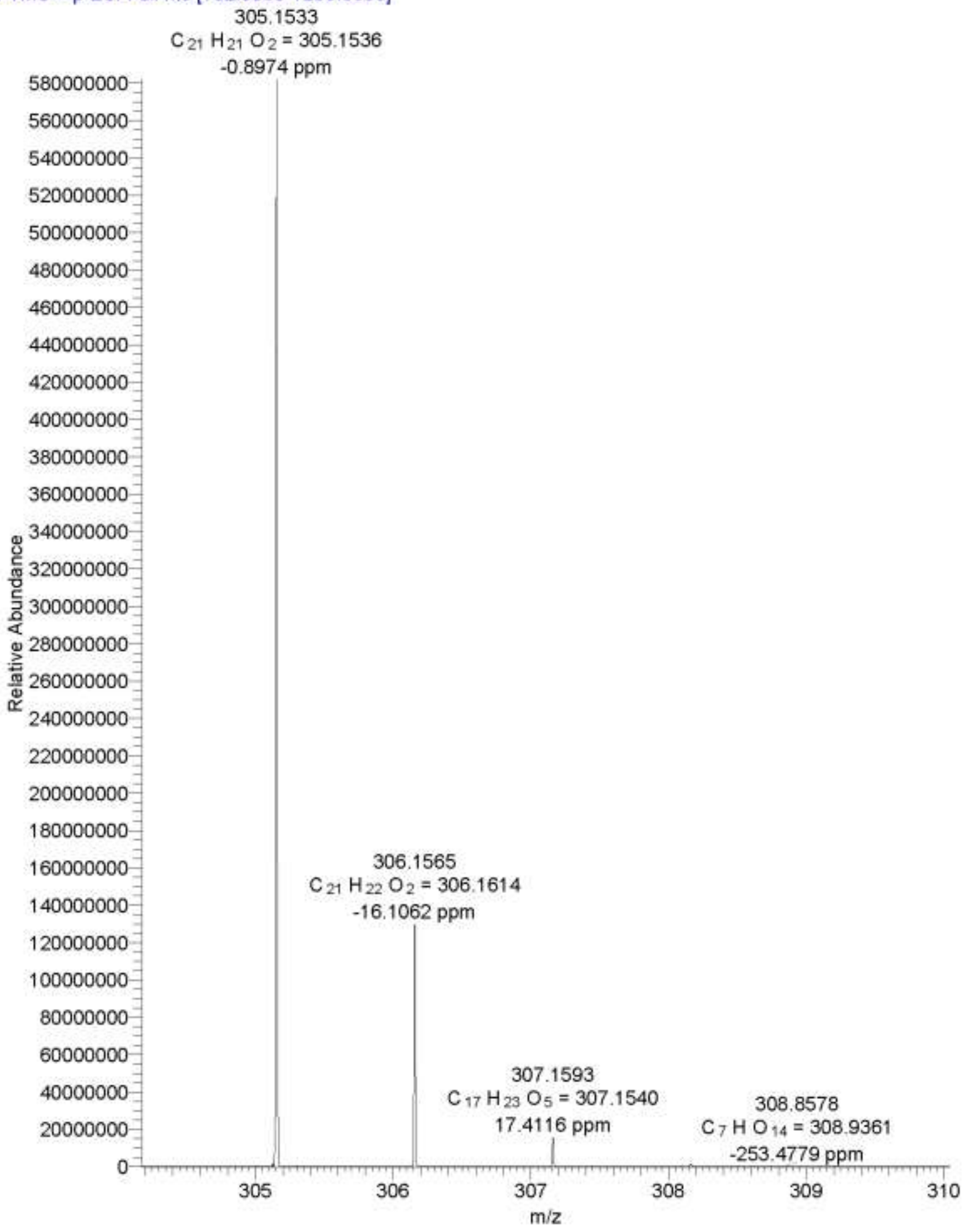

*ARM-3-124 126 127 ARM-3-126 28/07/2020 17.39.28*

Chromatogram

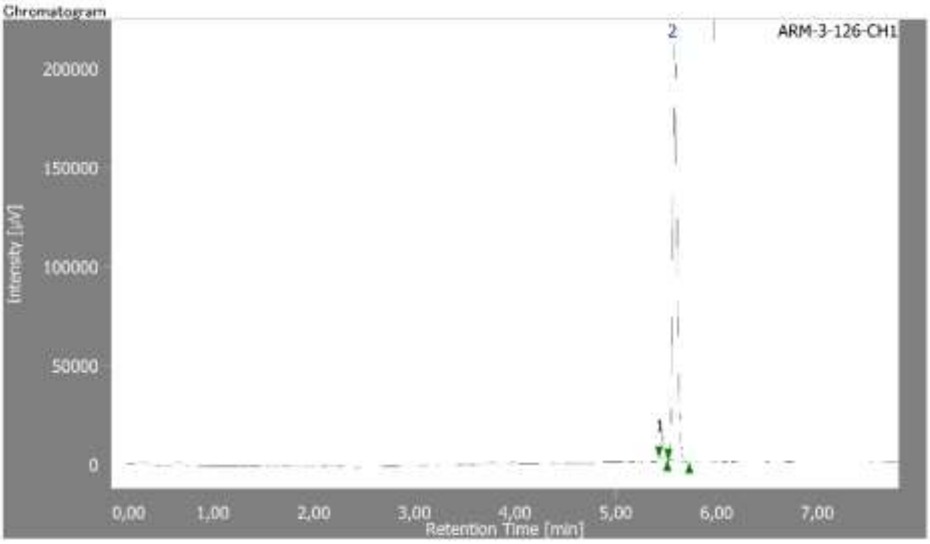


**Peak Information**

| # | Peak Name | CH | tR [min] | Area [µV·sec] | Height [µV] | Area% | Height% | Quantity | NTP | Resolution | Symmetry Factor | Warning |
|---|---|---|---|---|---|---|---|---|---|---|---|---|
| 1 | Unknown | 1 | 5.458 | 29297 | 11717 | 4.505 | 5.237 | N/A | 93883 | 1.697 | 1.238 | |
| 2 | Unknown | 1 | 5.583 | 621053 | 212040 | 95.495 | 94.763 | N/A | 85311 | N/A | 1.344 | |

Gradient Profiles - No data.

**ARM-3-124**

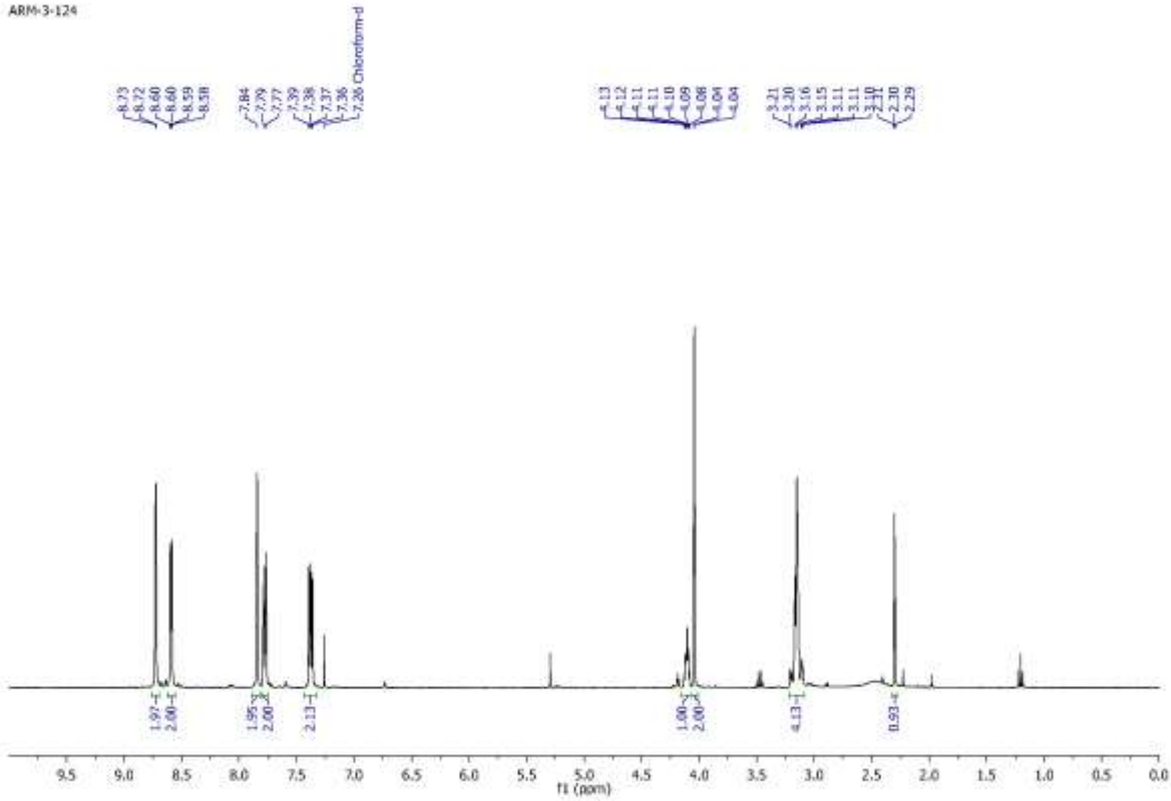
ARM-3-124
8.73
8.72
8.60
8.60
8.59
8.58
7.84
7.79
7.77
7.39
7.38
7.37
7.36
7.28 Chloroform-d
4.13
4.12
4.11
4.11
4.10
4.09
4.08
4.04
4.04
3.21
3.20
3.16
3.15
3.11
3.11
3.10
2.31
2.30
2.29
1.97
2.00
1.95
2.00
2.13
1.00
2.00
4.13
0.93
f1 (ppm)

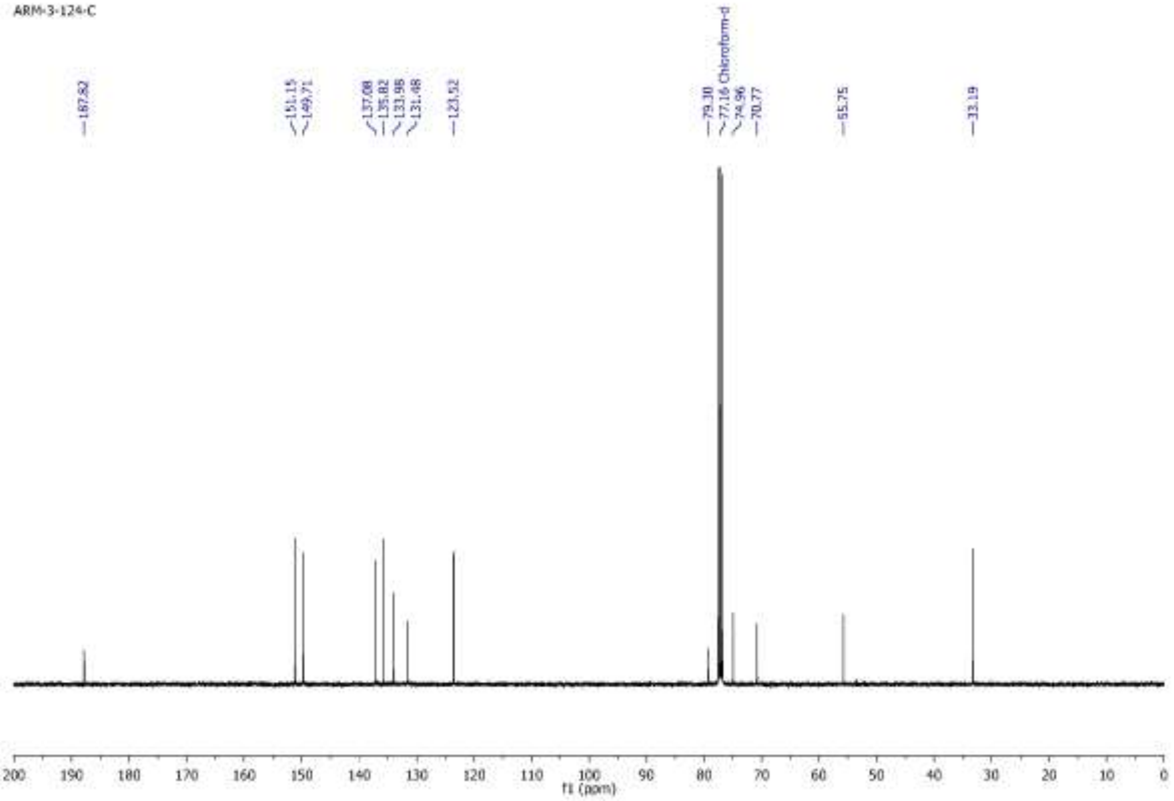
ARM-3-124-C
187.82
151.15
149.71
137.08
135.82
133.98
131.48
123.52
79.38
77.16 Chloroform-d
74.96
70.77
55.75
33.19
f1 (ppm)

ARM3-124 #682-717 RT: 5.39-5.65 AV: 18 NL: 4.52E9
T: FTMS + p ESI Full ms [132.0000-1200.0000]

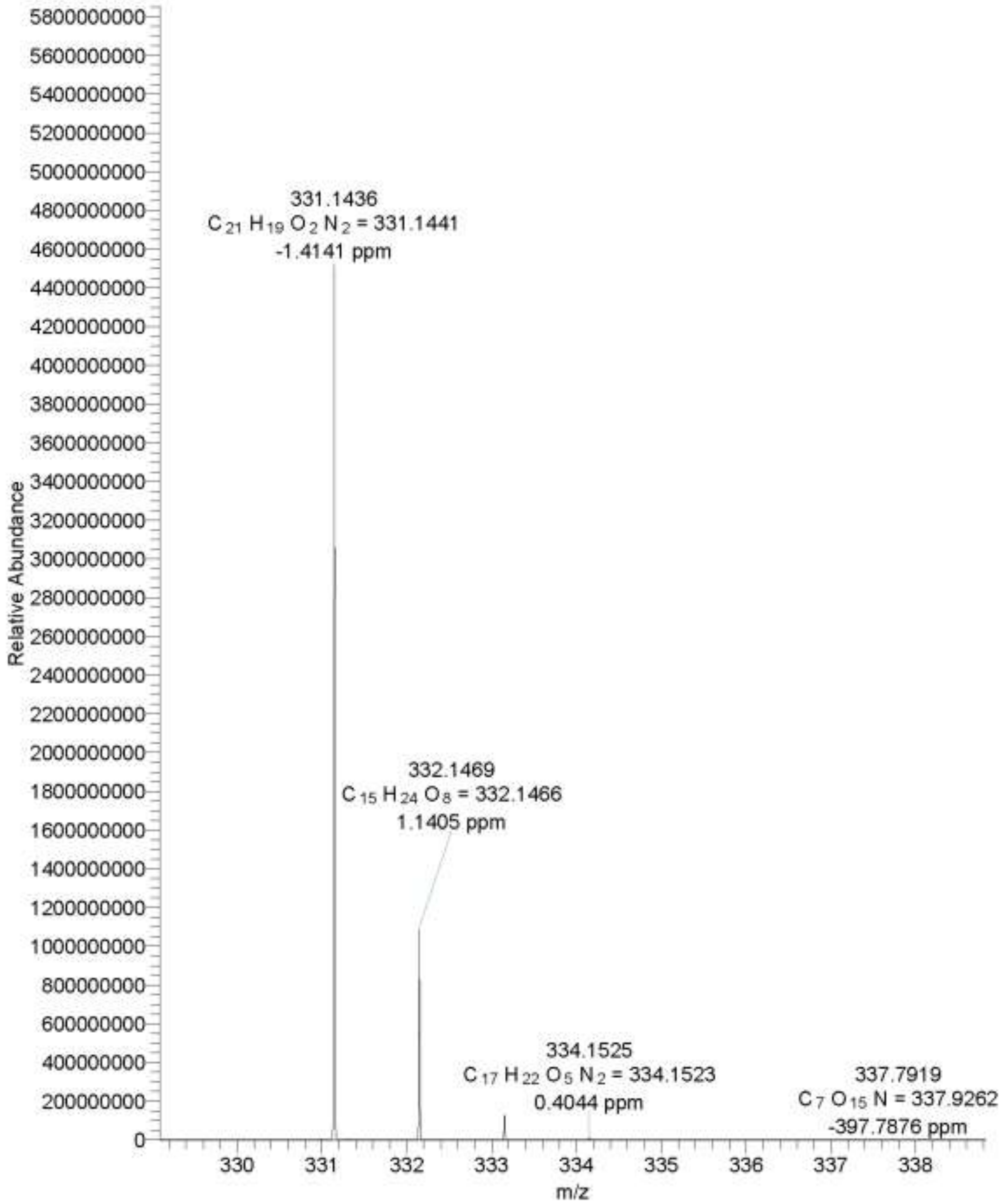

*ARM-3-124 ARM-3-124 23/07/2020 14:34:15*

Chromatogram

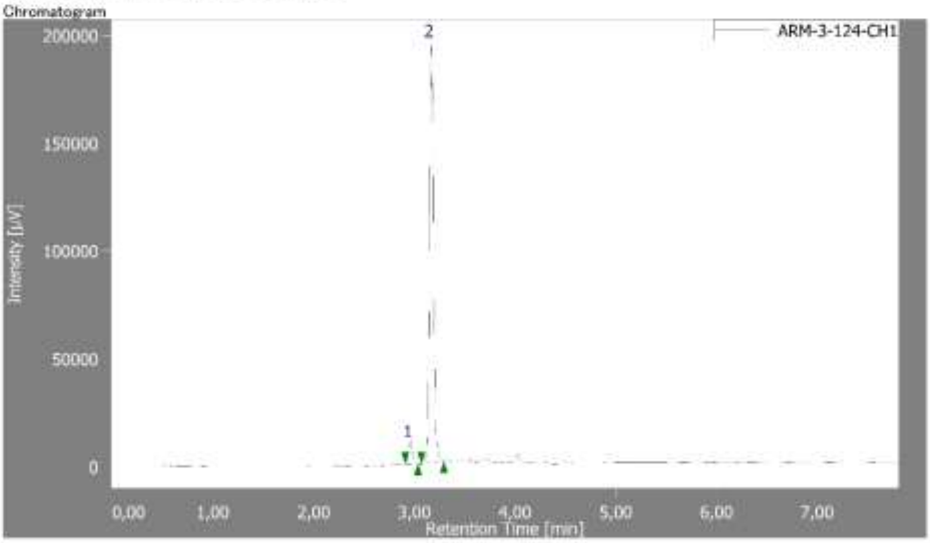


**Peak Information**

| # | Peak Name | CH | tR [min] | Area [µV·sec] | Height [µV] | Area% | Height% | Quantity | NTP | Resolution | Symmetry Factor | Warning |
|---|---|---|---|---|---|---|---|---|---|---|---|---|
| 1 | Unknown | 1 | 2.958 | 28214 | 10470 | 4.996 | 5.085 | N/A | 29487 | 3.027 | 1.138 | |
| 2 | Unknown | 1 | 3.167 | 536456 | 195422 | 95.004 | 94.915 | N/A | 33565 | N/A | 1.129 | |

**Control Method**

**ARM-3-127**

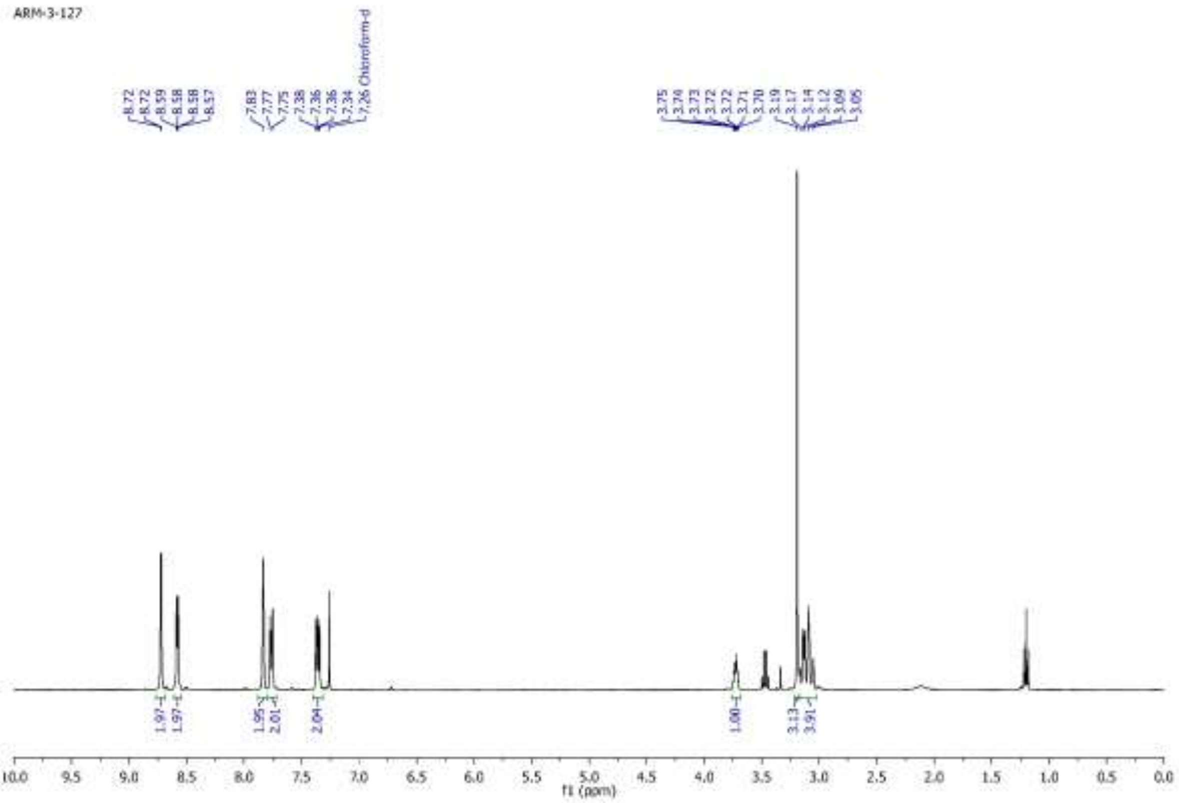
ARM-3-127
8.72
8.72
8.59
8.58
8.58
8.57
7.83
7.77
7.75
7.38
7.36
7.36
7.34
7.26 Chloroform-d
3.75
3.74
3.73
3.72
3.72
3.71
3.70
3.19
3.17
3.14
3.12
3.09
3.05
1.97
1.97
1.95
2.01
2.04
1.00
3.13
3.91
f1 (ppm)

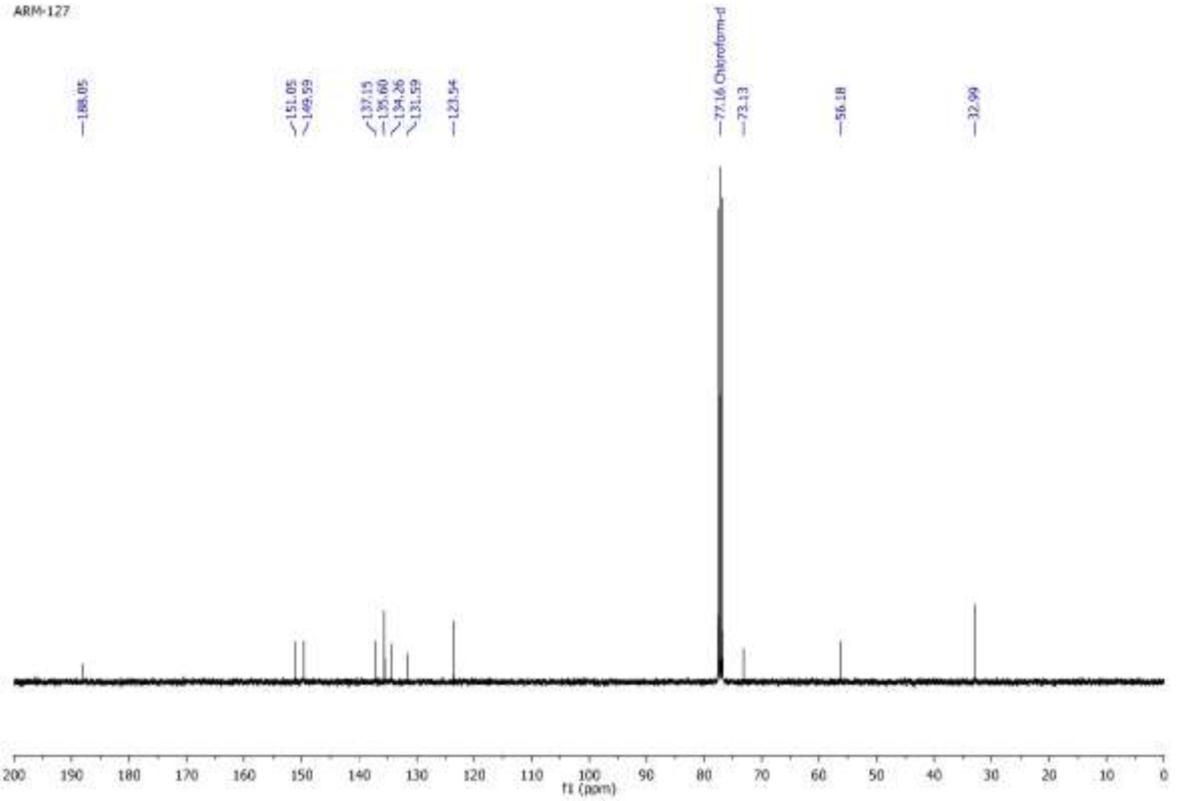
ARM-127
188.05
151.05
149.59
137.15
135.60
134.26
131.59
123.54
77.16 Chloroform-d
73.13
56.18
32.99
f1 (ppm)

C:\Users\...\Desktop\HRMS Mauro\ARM3-127 9/13/2020 9:12:09 AM

ARM3-127 #5-2786 RT: 0.04-21.98 AV: 1391 NL: 1.48E8
T: FTMS + p ESI Full ms [132.0000-1200.0000]

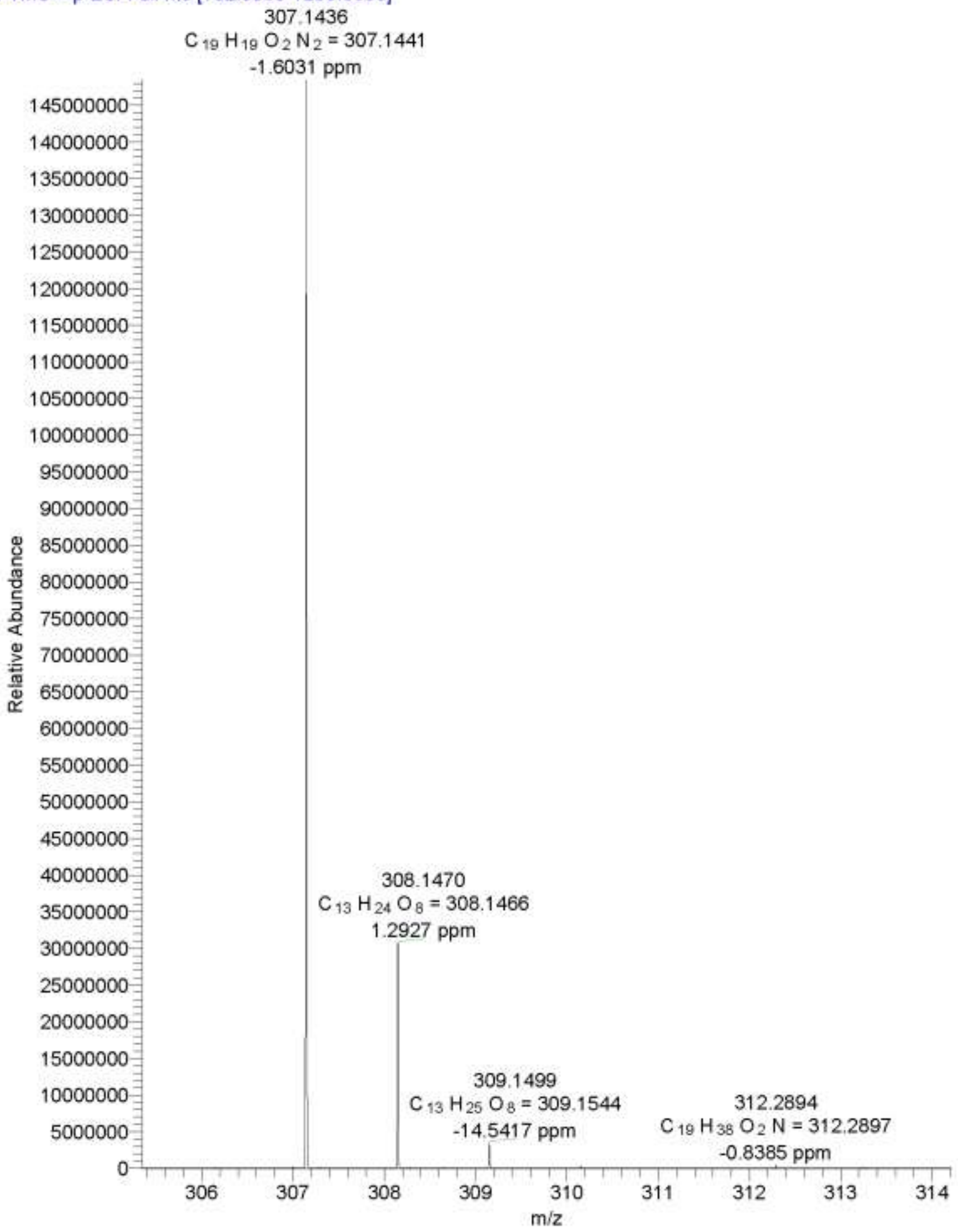

*ARM-3-124 126 127 ARM-3-127 28/07/2020 17:40:32*

Chromatogram

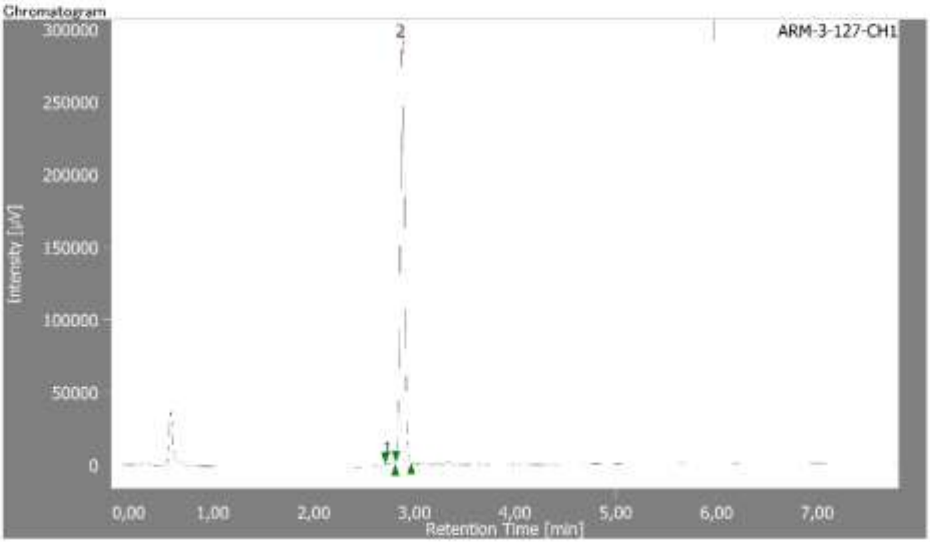


**Peak Information**

| # | Peak Name | CH | tR [min] | Area [µV·sec] | Height [µV] | Area% | Height% | Quantity | NTP | Resolution | Symmetry Factor | Warning |
|---|---|---|---|---|---|---|---|---|---|---|---|---|
| 1 | Unknown | 1 | 2.758 | 12335 | 4401 | 1.429 | 1.494 | N/A | 21378 | 1.628 | 0.898 | |
| 2 | Unknown | 1 | 2.883 | 850658 | 290090 | 98.571 | 98.306 | N/A | 21099 | N/A | 0.987 | |

Gradient Profiles – No data.
