## Supplementary Table 1 for "MCB-613 exploits a collateral sensitivity in drug resistant *EGFR*-mutant non-small cell lung cancer through covalent inhibition of KEAP1"

| | |
|---|---|
| Universal prefix | GGAAAGGACGAAACACCG |
| Universal suffix | GTTTTAGAGCTAGAAATAGCAAGTTAAAATAAGGC |
| Array_F | TAACTTGAAAGTATTTCGATTTCTTGGCTTTATATATCTTGTGGAAAGGACGAAACACCG |
| Array_R | ACTTTTTCAAGTTGATAACGGACTAGCCTTATTTTAACTTGCTATTTCTAGCTCTAAAAC |
| PCR1 F | GAGGGCCTATTTCCCATGATTC |
| PCR1 R | GTTGCGAAAAAGAACGTTCACGG |
| PCR2 F (i5 index) | AATGATACGGCGACCACCGAGATCTACAC(8-bp index)ACACTCTTTCCCTACACGACGCTCTTCCGATCT(0-5 bp stagger)TTGTGGAAAGGACGAAACACCG |
| PCR2 R (i7 index) | CAAGCAGAAGACGGCATACGAGAT(8-bp index)GTGACTGGAGTTCAGACGTGTGCTCTTCCGATCTACTTGCTATTTCTAGCTCTAAAAC |
| sgKEAP1_3 | GGAAAGGACGAAACACCGAATGAACACCATCCGAAGCGGTTTTAGAGCTAGAAATAGCAAGTTAAAATAAGGC |
| sgKEAP1_5 | GGAAAGGACGAAACACCGACAACCCCATGACCAATCAGGTTTTAGAGCTAGAAATAGCAAGTTAAAATAAGGC |
| KEAP1 F EcoRV | TAAGCAgatatcatgcagccagatcccagg |
| KEAP1 R NotI | TGCTTAgcggccgctcaacaggtacagttctg |
| KEAP1 1-60 F | ctggaggatcatacctagcaggcctttggca |
| KEAP1 1-60 R | tgccaaaggcctgctaggtatgatcctccag |
| KEAP1 1-178 F | ttcctggtgcagcagtaggaccccagcaatgc |
| KEAP1 1-178 R | gcattgctggggtcctactgctgcaccaggaa |
| KEAP1 1-314 F | gcacaagcccacgtaggtgatgccctg |
| KEAP1 1-314 R | cagggcatcacctacgtgggcttgtgc |
| KEAP1 1-597 F | gagcgaggtgacccgaatgtagtcgggccggagt |
| KEAP1 1-597 R | actccggcccgactacattcgggtcacctcgctc |
| sgNFE2L2_1 | GGAAAGGACGAAACACCGGCGACGGAAAGAGTATGAGCGTTTTAGAGCTAGAAATAGCAAGTTAAAATAAGGC |
| sgNFE2L2_2 | GGAAAGGACGAAACACCGTATTTGACTTCAGTCAGCGAGTTTTAGAGCTAGAAATAGCAAGTTAAAATAAGGC |
