## Supplementary Table 2 for "MCB-613 exploits a collateral sensitivity in drug resistant *EGFR*-mutant non-small cell lung cancer through covalent inhibition of KEAP1"

| | i7 index | i5 index | i5 stagger |
|---|---|---|---|
| PP1 | GAATTCGT | ATAGAGGC | a |
| PP2 | GAATTCGT | CCTATCCT | ca |
| PC9 1 | ATTACTCG | ATAGAGGC | a |
| PC9 2 | ATTACTCG | CCTATCCT | ca |
| PC9 3 | ATTACTCG | GGCTCTGA | gca |
| GR4 1 | TCCGGAGA | GGCTCTGA | gca |
| GR4 2 | TCCGGAGA | AGGCGAAG | tgca |
| GR4 3 | GAGATTCC | TATAGCCT | - |
| WZR12 1 | ATTCAGAA | TATAGCCT | - |
| WZR12 2 | ATTCAGAA | ATAGAGGC | a |
| WZR12 3 | ATTCAGAA | CCTATCCT | ca |
