## Supplementary Figure Legends for "MCB-613 exploits a collateral sensitivity in drug resistant *EGFR*-mutant non-small cell lung cancer through covalent inhibition of KEAP1"

### Supplementary Fig. 1. Diverse models of EGFR inhibitor resistance

a. Relative cell viability of parental PC9 cells, a gefitinib-resistant PC9 pooled population, and six gefitinib-resistant PC9 clonal populations following 72-hour incubation with gefitinib across an 8-point serial drug dilution series. Data are mean ± SEM for n = 3 biologically independent experiments.
b. Relative cell viability of parental HCC827 cells, a gefitinib-resistant HCC827 pooled population, and six gefitinib-resistant HCC827 clonal populations following 72-hour incubation with gefitinib across an 8-point serial drug dilution series. Data are mean ± SEM for n = 3 biologically independent experiments.
c. Relative cell viability of matched parental (MGH134) and osimertinib-resistant (MGH134 OR1 and OR2) patient-derived cell lines following 72-hour incubation with gefitinib across an 8-point serial drug dilution series. Data are mean ± SEM for n = 3 biologically independent experiments.

### Supplementary Fig. 2. QC and validation of CRISPR knockout screen

a. Heatmap depicting hierarchical clustering of plasmid pool duplicates, as well as initial ($T_0$) time points for each condition in singlicate and final (rep 1-3) time points for each condition in triplicate.
b. PCA plot depicting variance between plasmid pool duplicates, initial ($T_0$) time points in singlicate, and final (rep 1-3) time points for each condition in triplicate as explained by principal components 1 and 2.

c. Modified histogram depicting the variance explained by each principal component.
d. Immunoblot analysis of KEAP1 following CRISPR knockout with two sgRNAs targeting *KEAP1* versus a non-targeting control in parental PC9 and drug-resistant WZR12 cells.

**Supplementary Fig. 3. Validation of *NFE2L2* knockout**

a. Immunoblot analysis of NRF2 following CRISPR knockout with two sgRNAs targeting *NFE2L2* versus a non-targeting control in parental PC9 and drug-resistant WZR12 cells.
